# Emergent neutrality in consumer-resource dynamics

**DOI:** 10.1101/710541

**Authors:** Rafael D’Andrea, Theo Gibbs, James P. O’Dwyer

**Author notes:** **Statement of Authorship:** RD and JOD conceived the study. RD and TG performed analyses and wrote the paper. All authors contributed to the final version of the manuscript.

## Abstract

Neutral theory assumes all species and individuals in a community are ecologically equivalent. This controversial hypothesis has been tested across many taxonomic groups and environmental contexts, and successfully predicts species abundance distributions across multiple high-diversity communities. However, it has been critiqued for its failure to predict a broader range of community properties, particularly regarding community dynamics from generational to geological timescales. Moreover, it is unclear whether neutrality can ever be a true description of a community given the ubiquity of interspecific differences, which presumably lead to ecological inequivalences. Here we derive analytical predictions for when and why non-neutral communities of consumers and resources may present neutral-like outcomes, which we verify using numerical simulations. Our results, which span both static and dynamical community properties, demonstrate the limitations of summarizing distributions to detect non-neutrality, and provide a potential explanation for the successes of neutral theory as a description of macroecological pattern.

**Author Summary:** The neutral theory of biodiversity assumes that species are ecologically equivalent. Given the natural history observation of ubiquitous phenotypic differences between species, it is surprising that neutral theory has successfully predicted a broad range of biodiversity patterns, and simultaneously unsurprising that these results have not convinced ecologists that the natural world is neutral. However, we have lacked a description of how neutrality can emerge in a natural way from ecological mechanisms and species differences. Our study sheds light on this question, providing a theoretical backdrop for the success of neutral theory as a description of macroecological pattern. We derive a prediction for the degree to which consumers must differ in preferences for different resources before the resulting biodiversity patterns become distinguishable from neutrality. These predictions, which we confirm using simulations, show that neutral-like outcomes are possible even when resource requirements across consumers are very far from neutral. Our results can be tested in experimental microbial communities, where, equipped with an inferred consumption network, our analysis can yield predictions for biodiversity patterns and community turnover at different taxonomic levels.

## Introduction

One of the central questions in community ecology is how species interactions affect community structure and ecological dynamics, and conversely whether summarized representations of the latter can be used to make inferences about the former. Macroecological patterns such as species abundance distributions in particular have been extensively studied, and often used in inference approaches [1–3].

However, these inference approaches have been called into question, especially following intense debate over the merits of the neutral theory of biodiversity [1, 4–10]. Neutral theory assumes that all species and individuals in a community are ecologically equivalent, and therefore community assembly is a purely stochastic process, with differences in species abundances resulting from demographic noise [11, 12]. This hypothesis has been heavily tested across many taxonomic groups and environmental contexts [13–16], and despite its radical assumptions, has had some success in describing macroecological patterns. Prominent among these successes is the prediction of a broad range of species abundances, matching the functional form for the species abundance distribution observed in multiple high-diversity communities [12, 16–18]. On the other hand, the theory has been criticized on two fronts: first, it is unclear whether neutral theory could successfully predict a broader range of community properties, particularly regarding community dynamics from generational to geological timescales [19–27]. Second, it has been argued that the patterns that neutral theory successfully fits do not uniquely reflect the underlying ecological properties of the species involved; in particular, it is possible that summarized indices of community structure and dynamics may look neutral even when species are not ecologically equivalent [2, 27–31].

Some previous studies of neutral outcomes in non-neutral dynamics have focused on phenomenological models of species interactions, such as Lotka-Volterra competition [32, 33]. One drawback of this approach is that the influence of demographic noise relative to ecological selection can be tuned as a free parameter, rather than resulting endogenously from the interactions between species and the resources they compete for. A study of consumer-resource dynamics has shown that neutral behavior may result from metabolic tradeoffs guaranteeing ecological equivalence between consumers [34]. However, consumers vastly outnumber resources in those ecosystems, and therefore coexistence—and the accompanying neutrality—requires those tradeoffs to be exact. It remains an open question whether species competing for resources may appear neutral when their resource requirements are not fine tuned, and we still lack a quantitative prediction for how large niche differences between consumers must be in order for those differences to be reflected at the community scale.

Here we present a model of stochastic, non-neutral consumer-resource dynamics, with which we investigate the potential for neutral-like outcomes from two perspectives: snapshots of community structure, and dynamics over multiple timescales. Using a deterministic version of our model, we derive predictions for a threshold between neutral and non-neutral behavior, which we test against numerical simulations of the full model. Specifically, we hypothesize that when the timescales of relaxation to equilibrium under non-neutral dynamics are commensurate with the timescales of drift to extinction under neutral dynamics, the system will appear neutral. We investigate two qualitatively different departures from neutrality to test this hypothesis: a generalist scenario, where consumers have different preferences for resources but without a marked preferences for any one resource over others; and a specialist scenario, where each consumer has its own preferred resource, while consuming all other resources at a lower rate. Our analysis is based on these two particular ways to ‘break’ neutrality, but is not confined to them. However, we demonstrate that these cases support our hypotheses: consumer resource models with a degree of non-neutrality anywhere below our predicted threshold display neutral-like dynamics and patterns.

## Methods

### Consumer-Resource Model

We base our analysis on a stochastic model of *S* consumers, whose abundances we denote by *N*_*i*_ for *i* ∈ {1, …, *S*}, that compete for *K* abiotic, substitutable resources, whose concentrations are *R_k_* for *k* ∈ {1, …, *K*}. Resources are externally supplied, and depleted only by consumption. Consumers grow purely through the consumption of these resources, and also undergo density-dependent mortality. While this is a challenging model to solve exactly, we can straightforwardly simulate a stochastic process with these three event types (resource inflow, consumption, and consumer mortality). In this simulation approach, we define *T*{*X* → *X* + 1} as the transition rate at which the species with abundance *X* gains one individual, while all other abundances stay precisely the same. In this notation, the following transition rates represent the events that occur in our stochastic process:

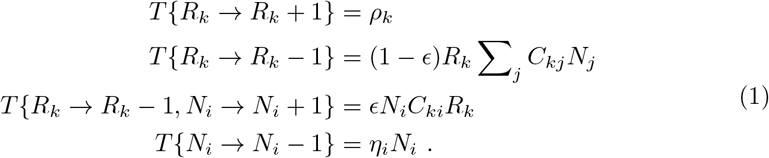

The basic event types that lead to the model in Eq 1 are conceptually represented in Fig 1. *ρ*_*k*_ is the inflow rate of the *k*-th resource. *C*_*ij*_ are non-negative coefficients which measure the rate of consumption of resource *i* by consumer *j*, forming a *K* × *S* matrix, *C*. In this study we set *K* = *S* for simplicity of the analysis, but our main mathematical results hold in general (see S1 Appendix section 3). *ϵ* measures the consumers’ efficiency in converting resources into consumer biomass, being the probability that a consumption event between resource *k* and consumer *i* results in a new individual of consumer *i* (in other words, every consumption event results in a loss of one resource unit, and a fraction *ϵ* of them also result in a gain of one consumer). We originally set *ϵ* to a small fraction, reflecting a need for many resource units to be consumed before each reproduction event, but this had no effect on outcomes other than to slow down the dynamics. Thus we set *ϵ* = 1 for expediency. *η*_*i*_ is the per capita mortality rate for the *i*-th consumer.

**Fig 1.**
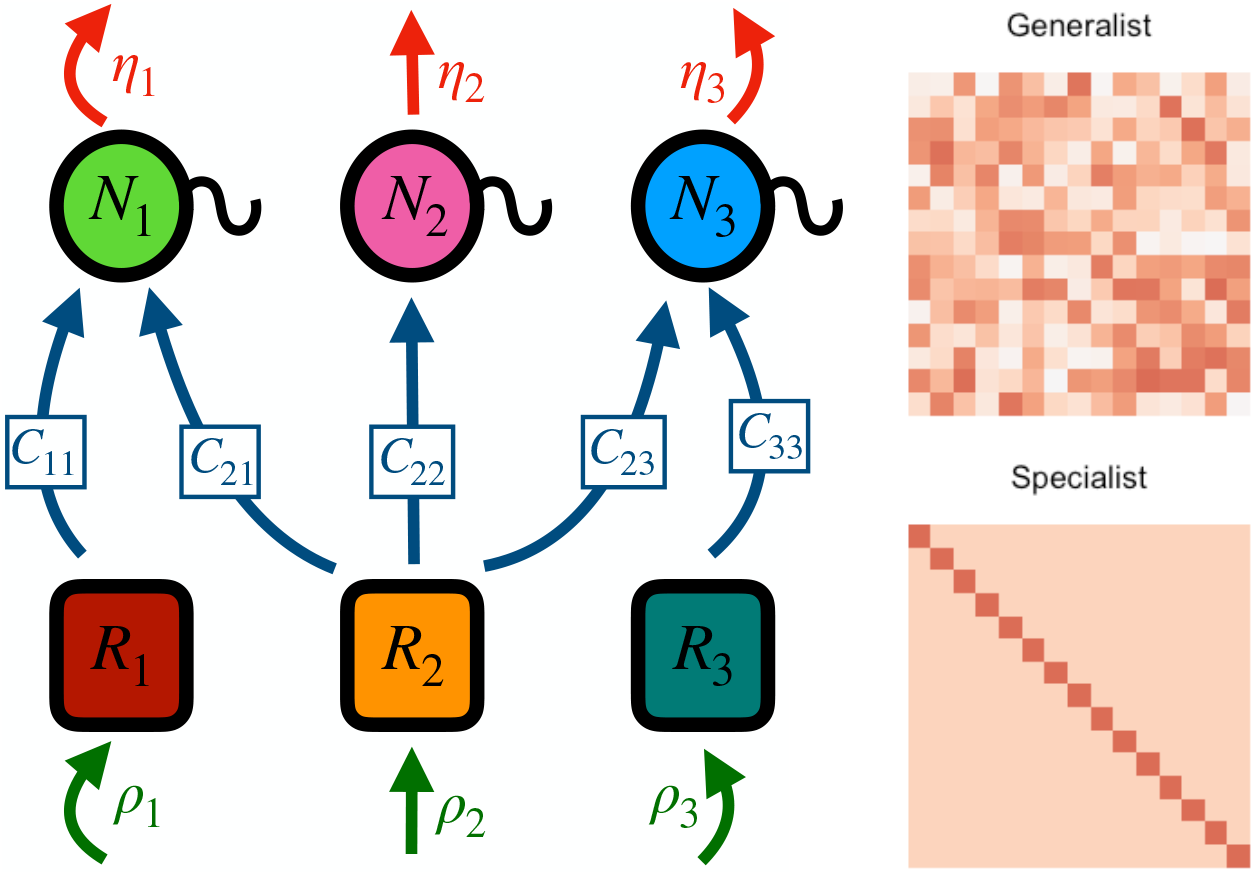
Model scheme. The left panel is a schematic of the models in Eqs 1 and 2. Resources flow into the system at fixed rate *ρ_i_*, while consumers die at rates *η_i_N_i_*. The arrows connecting resources to consumers represent non-zero consumption coefficients *C*_*ij*_. In this example, all consumers are capable of consuming resource 2, so there is community-wide competition for it. At the same time, consumer 1 is the only consumer that can utilize resource 1 and, similarly, consumer 3 is the only consumer that can deplete resource 3. The right panels are examples of the *C* matrices resulting from our two different parametrizations. In the generalist case, each entry is an independent sample from the same probability distribution with positive support. In the specialist case, consumer *i* can consumer resource *i* more quickly than any other resource. Other than this special resource, consumers consume every other resource at the same rate.

Every consumer goes extinct over long enough time scales, but in any real ecological system, speciation will tend to maintain species richness. Here we represent this balance in a simple way: every time a species with abundance *N*_*i*_ = 1 is selected for a death event (extinction), we keep it at *N*_*i*_ = 1, while logging the time it took for the death event to occur. This is akin to point speciation [35], although for simplicity we introduce the ‘new’ species with the same resource profile as the ‘extinct’ species. Alternatively, this process could be interpreted as very rare immigration events for that same species. While speciation and/or immigration can be modeled at increasing levels of realism (e.g. [36, 37]), our results are primarily based on the dynamics of extant species, and hence are largely insensitive to the details of this process. We confirmed this robustness for the specialists scenario under an alternative speciation/immigration scheme (Fig 10 in S1 Appendix).

In parallel, we consider a deterministic version of the same model:

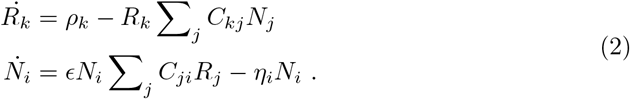

Using this deterministic model, we can calculate consumer and resource abundances at equilibrium. We will focus on coexistence solutions, i.e. equilibria of Eq 2 where all resources and consumers have positive equilibrium abundances [38]. While ecological disturbance or other perturbations can lead to important transient or cyclic behavior [39, 40], stable equilibria attract all nearby transient states, and therefore have a special role in determining the behavior of a dynamical system. Accordingly, they have been studied extensively in community ecology [41–43]. Let’s denote the abundances by vectors 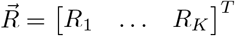 and 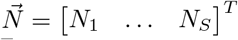 and the inflow and mortality rates by vectors 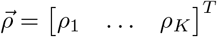 and 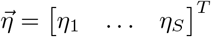. Then, given an invertible matrix *C*, we can choose positive inflow and mortality vectors 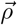 and 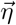 that lead to any positive abundance vectors (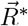 and 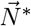) as equilibrium solutions of Eq 2. For simplicity, we will choose 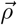 and 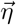 so that all resources converge to a single value *r* and all consumers converge to a value *n*. The values of the fixed points of Eq 2 correspond approximately to the average values of the abundances in Eq 1, so the stationary distribution for each consumer and resource will have a mean of *n* and *r* respectively.

In both Eq 1 and Eq 2, consumer *i* is determined by its consumption preferences (the *i*-th column of *C*) and mortality rate *η*_*i*_. Fixing a given set of equilibrium abundances determines 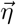, leaving the principal object of biological interest as the set of consumption preferences in *C*. For example, if all entries in *C* are identical, we recover a neutral limit where all consumer species have the same resource preferences. In this study we consider two different ways to move away from this neutral limit, shown in Fig 1. The first, which we call the specialist scenario, is when *C* has all diagonal entries equal to one value *C*_*d*_, while all off-diagonal entries are equal to another value *C*_*o*_, where *C*_*d*_ > *C*_*o*_. In this case, each consumer has a unique preferred resource that they consume more quickly than all others. Other than this special resource, each species consumes all other resources at the same rate. If *C*_*d*_ = *C*_*o*_, then all species are equivalent and we expect the stochastic version of this model to behave neutrally. In our second parametrization, called the generalist scenario, we sample each entry of *C* independently from a uniform distribution with mean *μ* and variance *σ*^2^. When *σ*^2^ = 0, the entries of *C* are simply *μ* and we expect the system to behave neutrally. As such, the magnitude of the ratio *C*_*d*_/*C*_*o*_ in the specialists scenario, and the coefficient of variation *CV* = *σ*/*μ* of the matrix in the generalists scenario quantify the system’s departure from neutral consumption preferences. Note that when *CV* = 0 in the generalist scenario and *C*_*d*_/*C*_*o*_ = 1 in the specialist scenario, the species abundance distribution (SAD) can be shown analytically to be a log-series (see S1 Appendix section 2).

In the next section, we predict the threshold values of these two measures of non-neutrality, below which we predict the community dynamics will look neutral. In other words, we predict how far from neutrality the model can be and still display neutral outcomes.

### Neutral-Niche Threshold

Species abundances in our stochastic process are constantly being perturbed away from the deterministic equilibrium by demographic noise. At the same time, consumer-resource feedbacks stabilize those abundances. The consumer community may therefore display neutral- or niche-like dynamics depending on the balance of influence between these forces. Because it is difficult to calculate the rates at which the abundances return to equilibrium in the stochastic model, we use the dynamics of the deterministic model near equilibrium as a proxy for the stabilizing force that the consumers experience in the stochastic system. In S1 Appendix, we show that the timescales of the dynamics of resources and consumers are naturally quite different, such that resources reach equilibrium and respond to perturbations much faster than consumers. As a result, resource abundances quickly converge to Poisson distributions (see Figs 3, 4 in S1 Appendix). This time-scale separation recapitulates the classic expectation of fast resource dynamics relative to consumer dynamics, but does not alone tell us whether or not the consumer system will be neutral-like. We thus turn our attention to consumer behavior for the remainder of this paper.

In order to identify a transition between niche and neutral dynamics, we consider a neutral community where the average species abundance is *n*. Let 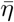 be the average consumer mortality rate (such that 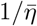 is the expected lifespan of an individual) and let *T_n_* be the expected time to extinction (measured in individual lifespans) for a species undergoing drift from its mean abundance. We then use 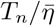 as a characteristic timescale for drift in our model—i.e. 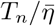 is the timescale over which drift takes a species with average abundance all the way to extinction. On the other hand, we use 1/|λ_+_| as the timescale for the stabilizing mechanism, where λ_+_ is the most negative eigenvalue corresponding to consumer dynamics in the deterministic model (i.e. λ_+_ determines the fastest timescale among the consumer dynamics). If the characteristic time for consumer abundances to deterministically return to equilibrium is much longer than the timescale for drift to cause large abundance fluctuations (ie. if 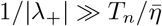), then drift should dominate consumer dynamics. This inequality also suggests a threshold at which drift ceases to be a good description of the abundance patterns, namely when 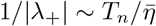.

We should note here that our estimate of the characteristic timescale of the stabilizing mechanism assumes that the linearized system is a good approximation to the actual dynamics throughout a large range of abundances. In fact, the linearized system is only guaranteed to be a good description in the neighborhood of the fixed point, but we hypothesize that it yields a good order-of-magnitude estimate for this transition more generally (for another example of this approach, see [44]).

In the S1 Appendix, we derive λ_+_ in both of our parametrizations of *C*, and from it obtain analytical expressions for when drift should be a good description of abundance patterns at equilibrium as a function of the other parameters in the model. For the generalist parametrization, we find that the threshold for the coefficient of variation *CV* = *σ*/*μ* is

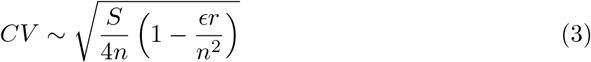

so the square of the threshold CV value should inversely depend on the mean consumer abundance *n*, if we disregard the second term in the parenthesis, which is small for our parameter choices. For the specialist parametrization, we derive a threshold scaling for the ratio between diagonal and off-diagonal elements of the consumption matrix, and find that

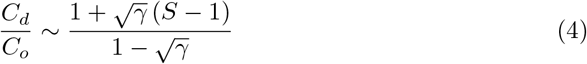

where 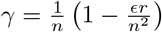. So, the specialist threshold is linear with the number of consumers in the community.

### Testing for Neutrality

We now test our predicted threshold by comparing our observations to neutral predictions using both static and dynamical properties of the community. Neutrality can be generally understood as symmetry at the species level, which corresponds to indifference in the fate of individuals upon changing species labels among populations [45]. This can be achieved in several ways [45–47]. Here we take the definition of [12], whereby neutrality occurs when all individuals have the same probability of death and birth events regardless of species identity and abundance. This definition serves our purpose of showing that non-neutral species may display neutral behavior at the community level when niche differentiation is not sufficiently large.

Under neutrality, species abundances follow a distribution which, in high-diversity communities, converges to Fisher’s log-series [48, 49]. The probability that a species has abundance *k* in this case is

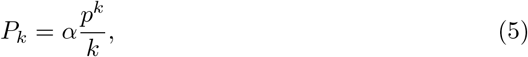

where *α* = −1/ log(1 − *p*), and the parameter *p* can be estimated from the mean abundance *n* by solving the equation *αp*/(1 − *p*) = *n*. We fit the log-series distribution to our simulated communities using the discrete Cramér-von Mises goodness-of-fit test [50], and consider the fit successful if the p-value exceeds 0.05.

To test our analytical predictions from the previous section, we define the neutrality threshold as the point of departure from neutral resource preferences at which the probability of a successful log-series fit drops below 50% (other choices for this probability cutoff do not change results qualitatively, see S1 Appendix). We determine this probability by fitting the log-series distribution onto ensembles of simulated communities, and running a logistic regression of the successful and rejected fits against the neutrality index of the different ensembles. In the generalist scenario, the neutrality index is the coefficient of variation in the consumption matrix, while in the specialist scenario it is the ratio between on- and off-diagonal entries.

We next define a test for whether long-timescale neutral dynamics fail to hold when the degree of non-neutrality of the consumer model passes through our predicted transition. Given a species current abundance *k*, its life expectancy (i.e. time to extinction) *T*_*k*_ under neutral dynamics has been shown to be [51]

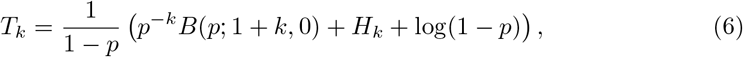

where *H*_*k*_ is the *k*-th harmonic number [52], *B*(*z*; *a, b*) is the incomplete beta function, and *p* is the log-series parameter (related to the speciation rate in neutral metacommunity models with speciation events). The extinction time in Eq 6 is given in generation units, with a generation time defined as the inverse of the mean mortality rate of consumers, 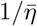, i.e. the expected lifetime of an individual. In our numerical simulations, we obtain life expectancy estimates as follows. Over the course of one simulation, after a burn-in period to ensure stationarity, we record all species abundances every 30 minutes of simulation time, corresponding to roughly 54 (212) generations in the generalists (specialists) scenario, for a total of 30,000 generations of simulation runtime in either scenario. An extinction event occurs when a species with abundance 1 is chosen for a death event. Because in our model the state *N*_*i*_ = 0 is not allowed, multiple extinction events may occur to the same species. Also, multiple abundance recordings may occur between successive extinction events, such that when an extinction eventually occurs, it will be linked to all previously recorded abundances since the last extinction. The time intervals between abundance recordings were sufficiently large to ensure independent data points, and the total runtime was long enough to observe extinctions in species with a wide range of initial abundances.

Finally, we test for neutral behavior at shorter timescales by comparing temporal fluctuations in species abundance against neutral predictions. For a subcritical stochastic birth-death process representing neutral dynamics, [51] derived an expression for the probability that a species has abundance *n* at time *t* given initial abundance *n*_0_ ([51], Eq. 2a). We use that expression to calculate the expected variance across histories of the stochastic process of the quantity 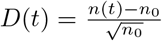[24], against which we compare our numerical observations. We use an ensemble of 100 histories, sampling the communities every generation for up to 100 generations. A species is considered to have zero abundance upon extinction, and we calculate the ensemble variance of *D*(*t*) across species with the same *n*_0_, and then average across all species with *n*_0_ ≥ 20 (the cutoff was necessary because low-*n*_0_ species displayed noisy behavior at longer timescales, and would require a larger ensemble).

## Results

In the generalist scenario, when the coefficient of variation (CV) in the consumption matrix is low (reflecting that consumers have similar preferences for all resources), the log-series distribution fits the SAD (Fig 2A). However, as the CV of the consumption matrix increases, the probability that the SAD conforms to the LS distribution declines (Fig 2B). For a community with 50 species and 50 resources, where the average species abundance is 245 and the average resource abundance is 100, a logistic regression indicates that the probability of a successful log-series fit drops below 50% when the CV of the consumption matrix is higher than 0.23. Defining this as the threshold for neutral-like patterns in the species abundance distribution, we note a power law between *CV* ^threshold^ and the mean species abundance *n* (Fig 2C), with an exponent close to our predicted value of −0.5 from Eq 3. A similar agreement occurs when we draw the consumption rates in the C matrix from a normal distribution rather than a uniform distribution (Fig 13 in S1 Appendix), as expected since the random-matrix theory we used to derive Eqns 3 and 4 is distribution-independent.

**Fig 2.**
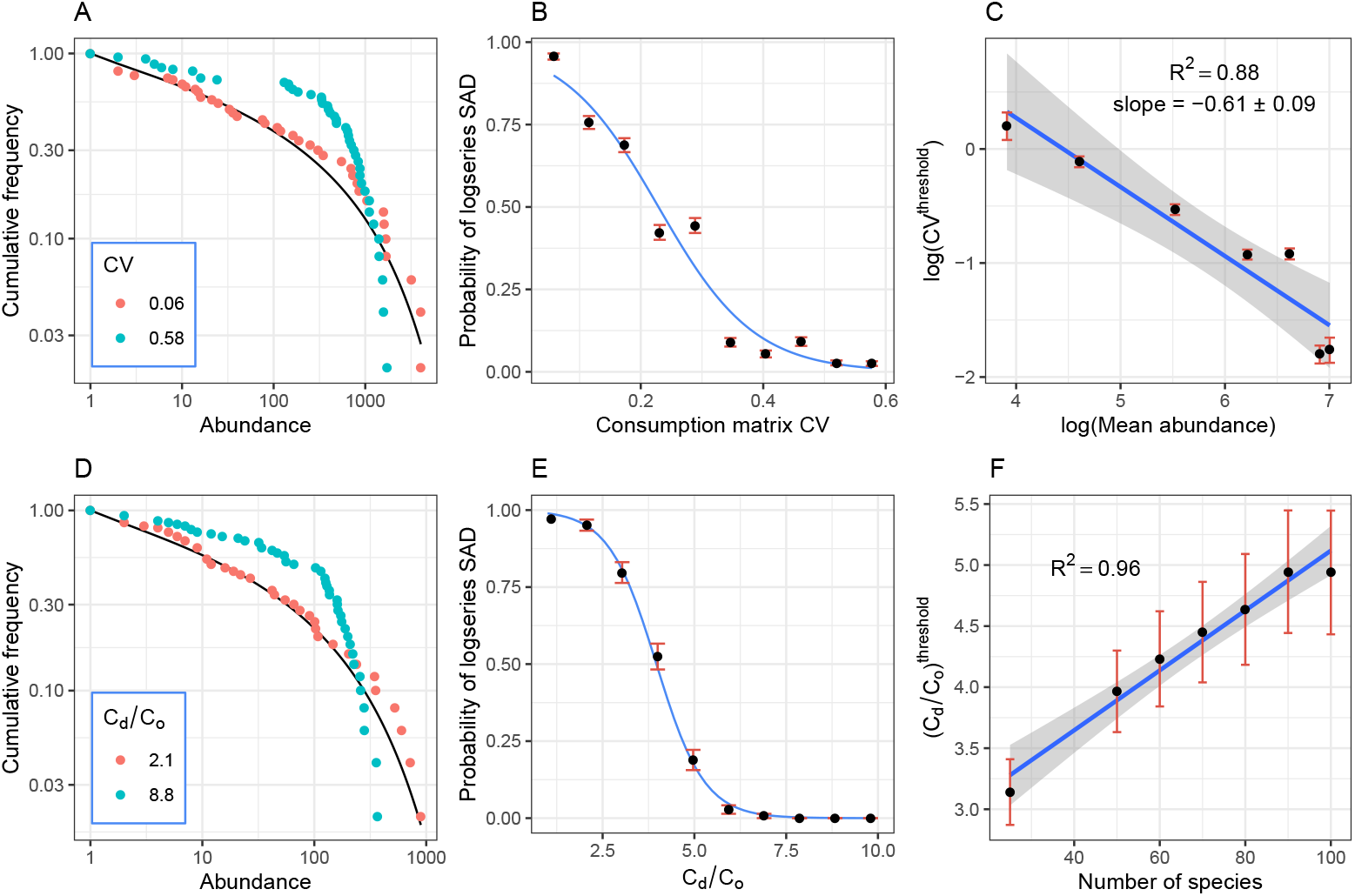
SAD results. Species abundance distribution (SAD) results for the generalists scenarios (A-C) and specialist scenarios (D-F). **A**: In the generalist scenario, the log-series distribution (black curve) fits the SAD when the coefficient of variation (CV) in the consumption matrix is sufficiently low (red points; Cramér-von Mises goodness-of-fit test p-value 0.78), but is rejected when the CV is sufficiently high (blue points; CvM test p-value 0.02). **B**: Probability that the logs-series distribution fits the SAD decreases with the CV of the consumption matrix. Points and error bars show the mean and standard error of the count of successful fits, out of an ensemble of 106 communities. Blue curve shows logistic regression. The threshold CV, defined as the point where the probability falls below 50%, is *CV* ^threshold^ = 0.23. **C**: log(*CV* ^threshold^) has a linear relationship with log(*n*), with slope −0.61 ± 0.09. This indicates a power law between *CV* ^threshold^ and *n*, with an exponent close to our analytic prediction of −0.5. Error bars show uncertainty propagated from the standard errors of the fitted parameters in the respective logistic regressions. Bands show the 95% CI of the linear regression. **D**: In the specialists scenario, communities with low *C*_*d*_/*C*_*o*_ ratio (red points; CvM test p-value 0.267) conform to the log-series distribution, while communities with sufficiently high *C*_*d*_/*C*_*o*_ (blue points; CvM test p-value < 0.001) reject the LS distribution. **E**: The probability of the LS distribution fitting the SAD decreases as *C*_*d*_/*C*_*o*_ increases, with the threshold at 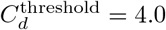 (we set *C*_*o*_ = 1). Each data point summarizes an ensemble of 143 communities. **F**: 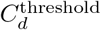 increases linearly with the number of species in the community, in qualitative agreement with Eq 4. Parameters: A-C: *K* = *S* = 50, *n* = 500, *r* = 100. D-F: *K* = *S* = 50, *n* = *r* = 100.

Results were analogous in the specialists scenario. Communities with low *C*_*d*_/*C*_*o*_ ratio, indicating a small-magnitude difference between preferences for main and secondary resources, conform to the log-series distribution, while communities with sufficiently high *C*_*d*_/*C*_*o*_ reject it (Fig 2D). Indeed, the probability that the log-series distribution fits the SAD decreases as *C*_*d*_/*C*_*o*_ increases (Fig 2E). In communities with 50 species and 50 resources, with mean species and resource abundances at 100, the threshold ratio for a successful log-series fit is 4.0. This threshold increases linearly with the number of species in the community (Fig 2F), as predicted by Eq 4.

Note we cannot use Eqn (4) to predict the slope in Fig 2F, as the formula for the threshold is defined up to a multiplicative constant. The reason for this constant is that we can estimate a characteristic timescale for relaxation after a perturbation under stabilized dynamics, but the exact time to relaxation depends on the size of the perturbation. We can always estimate the time it takes to get some arbitrary percentage of the way back to equilibrium–for example, the half-life of the decay back to the fixed point, but this introduces a multiplicative constant on the left side of our Eqns (3) and (4). The results shown herein are as quantitative as our formulas allow.

To compare the propensity for neutral-like abundance pattern across the generalist and specialist scenarios, we use a non-neutrality index applicable to both, based on the average cosine between the vectors representing the resource preference profiles of different species (i.e. the columns of the consumption matrix). The average cosine across all species pairs represents the average similarity between resource preference profiles. We therefore define the non-neutrality index as NNI = 1 − cos. The limit of complete neutrality, i.e. identical resource preferences among all species, corresponds to a cosine of 1, and thus NNI = 0. The opposite limit of complete niche differentiation, i.e. where each species consumes a single resource with zero overlap, corresponds to a cosine of 0 (NNI = 1). Fig 3 shows that in both the generalist and specialist scenarios, the probability of neutral-like abundance distribution is close to 100% in the neutral limit, as expected, but remains positive as we deviate from neutrality. The probability of rejecting the log-series only falls below 50% when the NNI is as high as 0.15 ± 0.05 in the specialist scenario, and 0.22 ± 0.03 in the generalist scenario (Fig 3). For a given NNI, the abundance distribution in the generalist scenario typically appears more neutral-like than in the specialist scenario.

**Fig 3.**
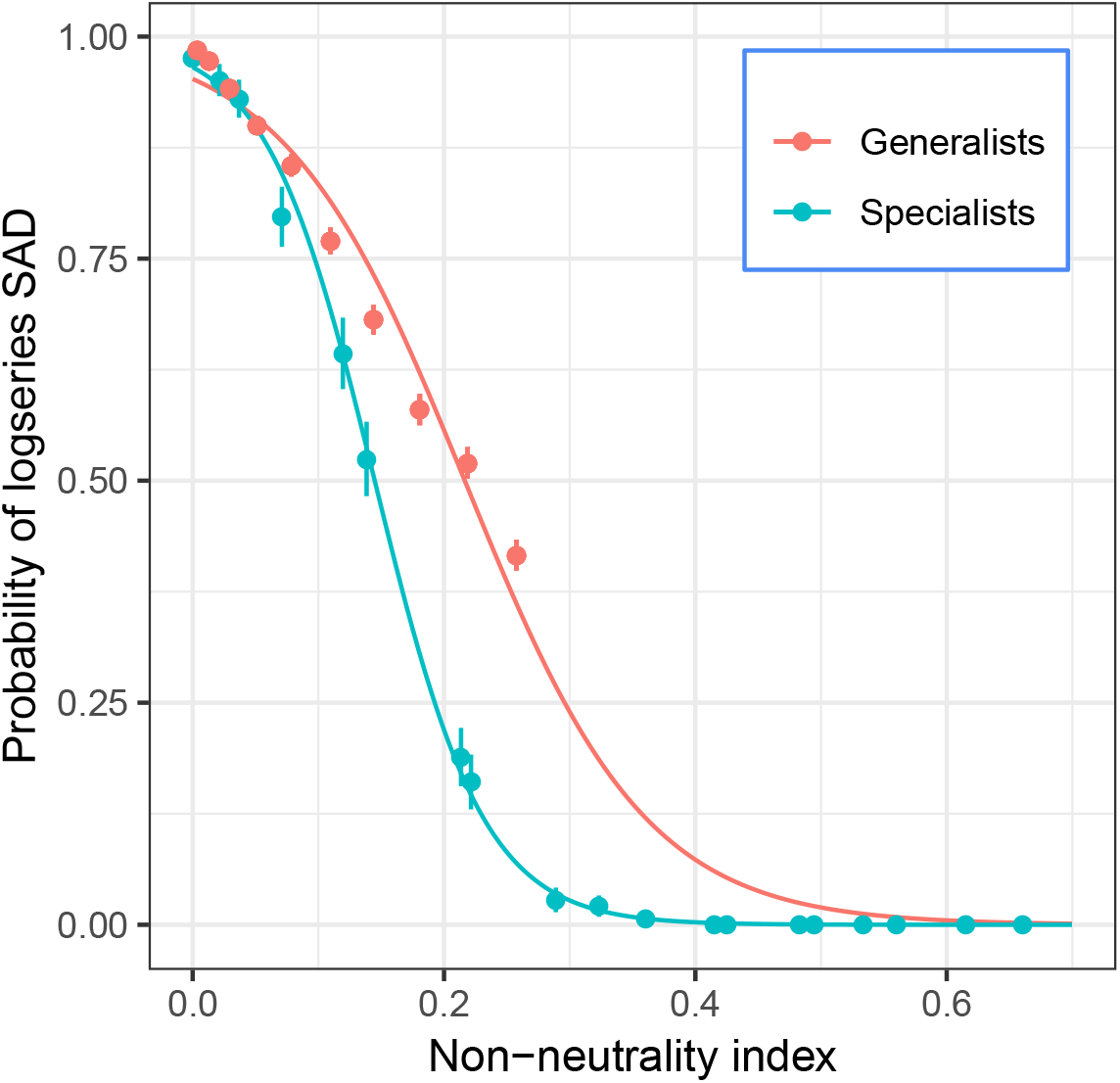
SAD results: generalists vs specialists. Probability of log-series SAD plotted against a non-neutrality index defined as NNI = 1 − cos, where cos is the cosine between vectors representing species resource preferences, averaged across all species pairs in the community. Complete neutrality would correspond to NNI = 0 (i.e. cos = 1), reflecting full overlap in resource preferences. For the same NNI value, communities in the generalist scenario are typically more likely to conform to the log-series distribution than communities in the specialist scenario. Parameters: *S* = *K* = 50, *n* = *r* = 100.

Extinction time results are shown in Fig 4. As expected, life expectancy increases with species abundance. In the generalist scenario, species extinction times in communities with low CV closely match predictions from the neutral model, whereas communities with sufficiently high CV depart from neutral predictions (Fig 4A). When there is a poor match, the neutral model underpredicts the extinction times, especially for species with high abundance (Fig 4A). Plotting observed extinction times against neutral predictions in communities with different CV reveals that higher CVs lead to increasingly poor matches to neutrality, especially for high-abundance species (Fig 4B). Interestingly, observed extinction times seem linearly related to predictions regardless of the CV. The slope of this relationship increases with the CV (Fig 4C), being close to 1 at low CV, indicating a perfect match to neutral predictions, and > 1 at higher CV. This indicates that species of high mean abundance have particularly long life expectancy beyond neutral expectations, suggesting that niche differentiation has a disproportionate stabilizing effect on common species.

**Fig 4.**
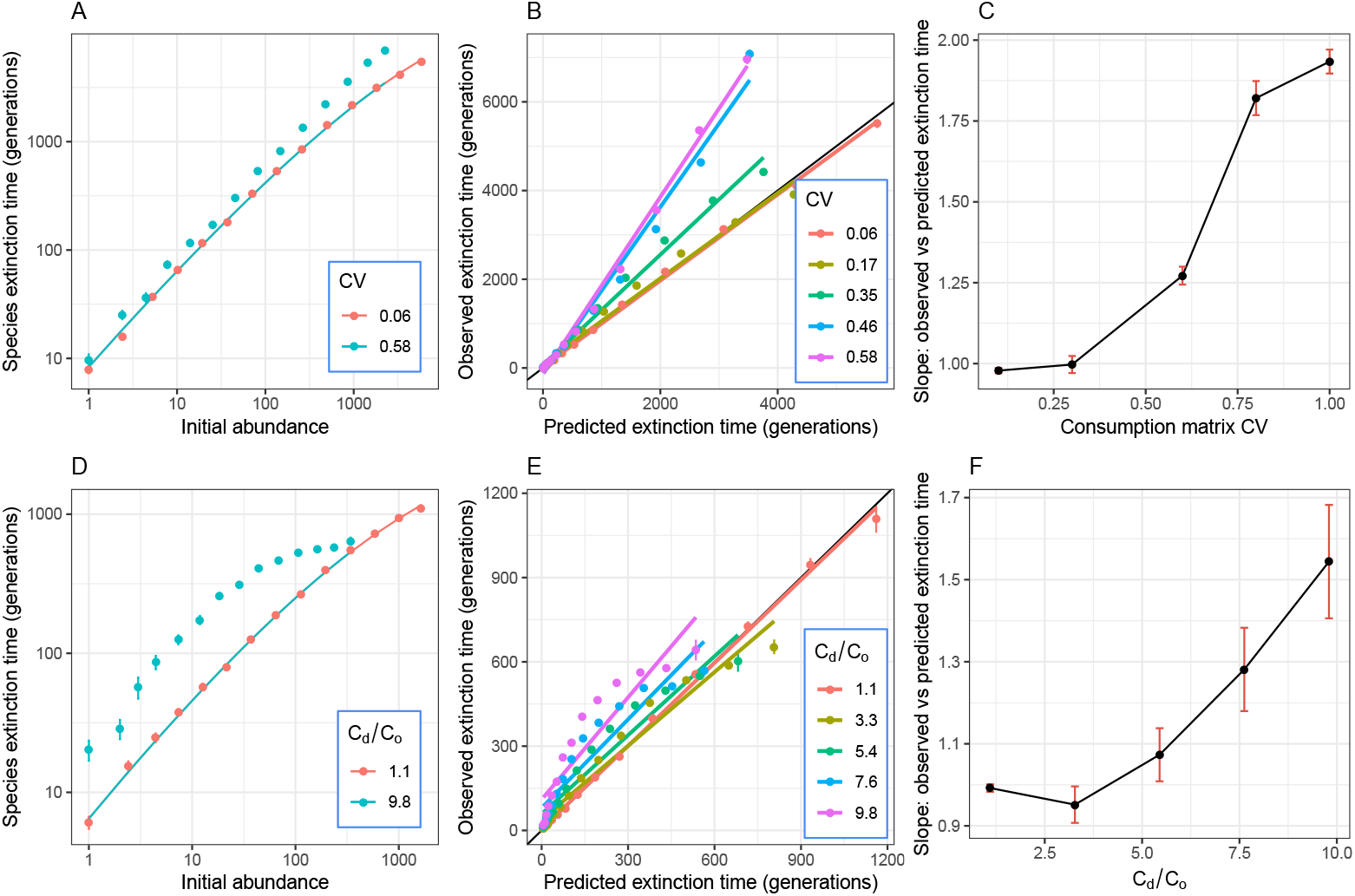
Extinction time results. Extinction time results for the generalist (A-C) and specialist (D-F) scenarios. **A**: Points and error bars show average and standard errors of extinction times for species in logarithmically binned abundance categories. Curves show neutral predictions. In the generalist scenario, species extinction times match predictions from the neutral model in communities with low CV, but consistently exceed neutral predictions in communities with high CV, especially for high-abundance species. **B**: Plotting observed versus predicted extinction times in communities with different CV (colors) reveals that those with low CV conform closely to the neutral predictions (black line illustrates a perfect match), while higher CVs lead to increasingly poor matches to neutrality, especially for high-abundance species. Note that extinction times seem linearly related to predictions regardless of the CV. **C**: The slope of this relationship increases with the CV, being close to 1 (perfect match to neutral predictions) at low CV and > 1 at higher CV. **D-F**: Results in the specialist scenarios are analogous to the generalist scenarios, except that extinction times of high-abundance species saturate. Parameters: A-C: *K* = *S* = 50, *n* = 500, *r* = 100. D-F: *K* = *S* = 50, *n* = *r* = 100. Summary statistics were obtained from ca. 5,000 to 20,000 data points for each abundance bin in the generalists scenario, and 1,000 to 7,000 data points in the specialists scenario.

The specialist scenarios showed analogous results to the generalist scenarios regarding matches to neutrality or lack thereof, except that extinction times of high-abundance species saturate. This could be because the restoring force in this scenario increases very quickly for large deviations from equilibrium, so a species that fluctuates to high abundance almost immediately returns to its mean abundance, thus not significantly increasing its life expectancy. By contrast, the generalist case lacks a strong stabilizing force, so excursions towards high abundance tend to substantially increase time to extinction.

Non-neutral community dynamics also displayed neutral-like behavior at shorter timescales (Fig 5). Temporal fluctuations in species abundances in the specialist scenario were indistinguishable from neutrality for up to 10 generations, even when the non-neutrality index was as high as 0.5. At longer timescales, niche stabilization tended to reduce the ensemble variance of 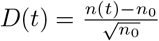 relative to neutrality, as expected. However, it was only at maximum non-neutrality (NNI = 1), corresponding to fully specialized and therefore non-interacting species, that var(D) was immediately distinguishable from neutral, plateauing within a few generations.

**Fig 5.**
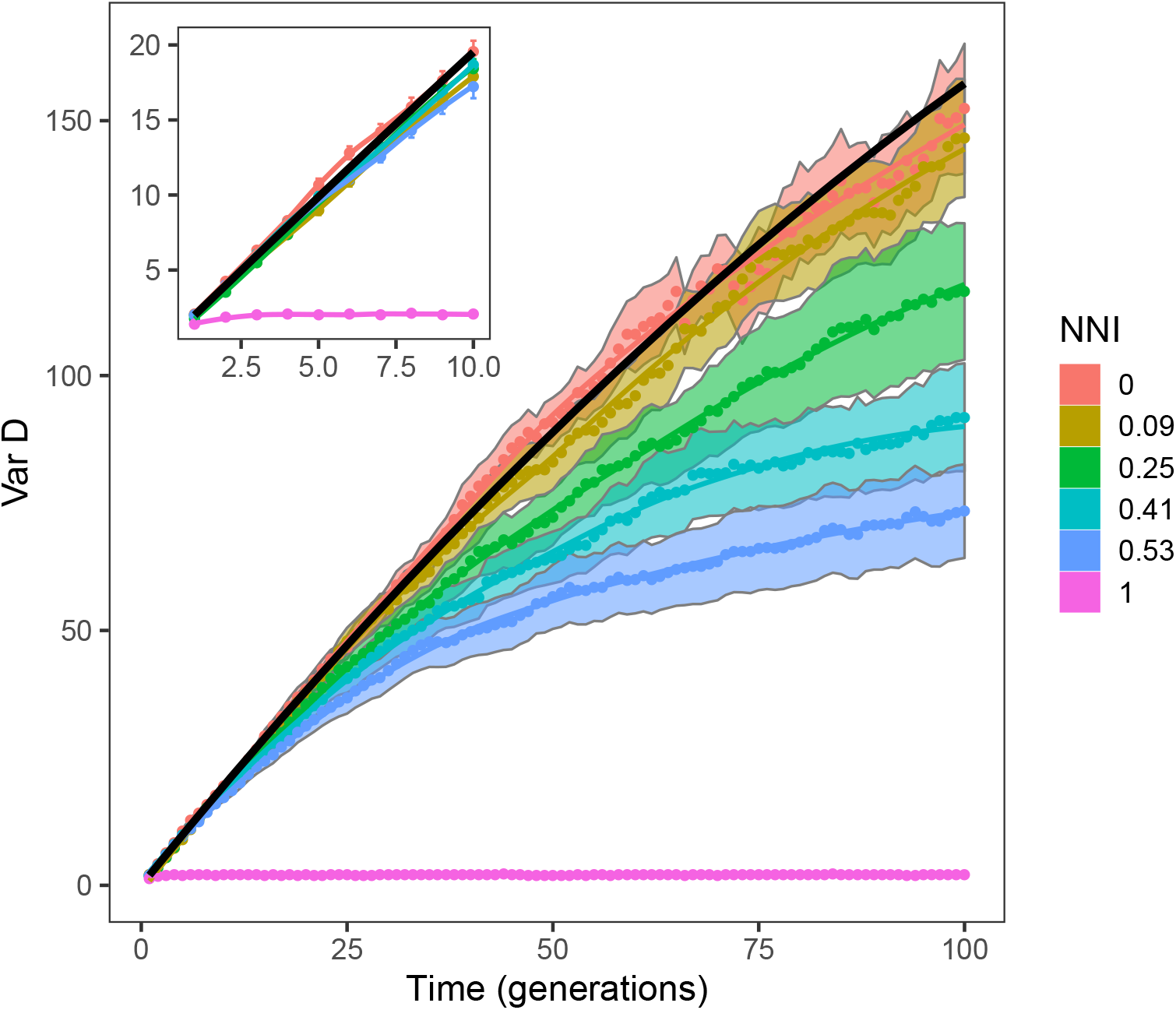
Shorter timescale results. Variance in species abundances over time, for different parametrizations of the specialist scenario. Vertical axis plots the variance across histories of the stochastic process of 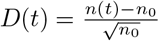, which is then averaged across species with different initial abundances *n*_0_ (bands show standard error of the mean). Colors show parametrizations with increasing non-neutrality index (NNI), with lines showing the loess regression with smoothing parameter set to 1. Black line shows neutral prediction. Inset highlights similarity of all curves except NNI = 1 at timescales up to 10 generations.

Notably, the neutral-like behavior observed here applies to communities whose resource preference profiles would be easily distinguishable from true neutrality, if they were directly measured. In other words, our predictions and results are not solely for “small” departures from neutrality, and yet we still observe neutral-like static and temporal patterns. To quantify this, we consider a neutral community of same richness and size as in Fig 3. If all consumption events under true neutral dynamics are observed over the course of an average species lifetime (in this case, approximately 250 generations), the 95% confidence interval for the average pairwise cosine is estimated to lie between 0.992 and 0.996 (see Fig 9 in S1 Appendix), well above the threshold for 50% rejection of the log-series abundance distribution. This means communities that are unmistakably non-neutral under direct observations of consumption events would still look neutral under summary distributions like the SAD or extinction times. Of course, in most if not all cases it is unfeasible to directly observe consumption events, especially over such long timescales. This impracticality is one reason why ecologists have indeed tested theories of community assembly using summarizing distributions.

## Discussion

We showed that communities of competing species may appear neutral despite differences in resource preferences that confer niche stabilization, and this behavior is quite predictable from summarized properties of the consumer-resource network. We provided quantitative underpinnings for why non-neutral dynamics may lead to neutral outcomes, namely because the niche stabilization provided by species differences in resource use profiles may not be strong enough to overcome the influence of demographic stochasticity in generating patterns at the community scale. Formally, this is reflected in the small eigenvalues of the Jacobian of the deterministic counterpart of our stochastic resource-consumer model.

Our simulation results qualitatively supported our hypothesis that when the relaxation time to equilibrium in the stabilized resource-consumer system is comparable to the timescales of drift to extinction under neutral dynamics, one cannot easily reject the log-series abundance distribution, and species life expectancies match neutral predictions. Furthermore, at generational timescales, even full knowledge of species abundances over time may not suffice to reject neutrality. This has implications for inferring niche differentiation in real life systems, such as distinct metabolic syndromes in microbial communities, where the differences between consumers may not be large enough to overcome the influence of drift.

Our study demonstrates the disconnect between neutrality at the pattern level—i.e. whether the distribution and temporal variation in species abundances are distinguishable from neutral predictions, and neutrality at the process level—i.e. whether species differ in resource preferences. In doing so, it builds on the point made by others that neutral SADs do not imply neutral ecology [1, 2]. Furthermore, our connecting emergent neutrality to stabilization timescales in a stochastic world provides a mechanistic explanation for the success of neutrality as a macroecological theory.

Earlier investigations of neutral-like outcomes in non-neutral communities have focused on phenomenological models of species interactions such as Lotka-Volterra competition [32, 33]. Our approach differs from these studies in two main ways. First, these approaches compare neutral dynamics to niche dynamics combined with exogenous immigration. That makes it challenging to isolate the stabilizing effects due to niche differentiation from those arising from immigration. By contrast, we model an essentially closed system and thus compare drift dynamics to stabilization driven almost purely by niche differentiation (strictly speaking, immigration is present but very low, only affecting the abundances of the rarest species). Second, in these stochastic Lotka-Volterra approaches, the corresponding deterministic dynamics do not completely determine the stochastic transition rates, leaving the degree of demographic noise as a free parameter that can be tuned to adjust the balance of noise relative to deterministic ecological selection. In our stochastic consumer-resource model, the level of noise is set endogenously by the structure of the Gillespie simulation. Our finding that neutral dynamics are still possible demonstrates that drift is a natural outcome of self-contained niche-differentiated ecological dynamics without tuning the degree of noise. While the previous approaches analytically solve simplified versions of stochastic Lotka-Volterra dynamics, we derive an analytical prediction for when and why the full stochastic consumer-resource model appears neutral.

In this context of consumer-resource dynamics, neutral behavior has been previously shown to arise from consumers competing for a small number of resources [34]. In that study, consumers satisfy fine-tuned metabolic tradeoffs that allow an arbitrarily large number of species to coexist on a neutrally stable manifold of fixed points, on which drift fully drives the stationary distribution. However, if consumers violate the precise metabolic tradeoff constraints, then only a small number of consumers coexist at one fixed point, and the stationary abundance distribution of these consumers is no longer guaranteed to appear neutral. In contrast, the numbers of consumers and resources are equal in our modeling framework, so the consumers coexist at a unique fixed point. In our stochastic model, it is the timescale of the stabilizing dynamics, which is determined by the degree of heterogeneity in resource preferences, that gives rise to neutral dynamics.

The neutrality threshold herein defined is a useful construct for testing our analytical predictions, not a methodological prescription for field ecologists. Indeed, the threshold’s numerical value depends on the choice of cutoff for the probability of rejecting the log-series. In reality, this probability increases gradually rather than abruptly with increasing departures from neutral resource preferences. Furthermore, one’s ability to reject neutral pattern in a community of interest depends not only on similarities between neutral and non-neutral outcomes, but also on the statistical power of goodness-of-fit tests. As such, in order to test our analytical predictions, the threshold’s numerical value is less meaningful than how it scales with changes in parameters such as community size and richness. We verified that this scaling is robust to different cutoff choices, and also to different tests and methodological approaches for rejecting neutrality (see Fig 8 and Fig 11 in S1 Appendix).

In nature, the consumption matrix could be far more complex than the two cases considered here, reflecting diverse metabolic strategies of consumers from different environments and evolutionary histories. For example, groups of the columns of *C* could be highly correlated when the corresponding groups of consumers perform similar functional roles in the community. The community as a whole could also exhibit a stronger preference for certain resources over others, introducing correlations between the rows of *C*. We ignored these complications for the sake of analytical tractability, but our results are likely generalizable: it is the bipartite structure of consumer-resource models, and not the specific structure of *C*, that separates the spectrum of our deterministic model into consumer and resource bulks. Because it is this separation that causes long excursions from the mean abundance in the stochastic model, our results suggest that a transition from neutral to niche dynamics will be a feature of consumer-resource models with more complex consumption preferences than considered here.

The fact that abundances in specialist communities are easier to distinguish from neutrality than in generalist communities with similar niche overlap suggests that specializing towards a single resource has a stronger niche-like impact on abundance distributions than unstructured variation in resource preferences. It follows that departures from neutrality are not all equal with regards to impact on abundances, leaving open the possibility that some special structure in the consumption matrix may very quickly lead to detectable differences from neutral pattern. One method to test this possibility would be to infer the consumption matrix by fitting our consumer-resource model to experimental abundance measurements from monoculture experiments on each resource. Then, different consumption matrix structures could be investigated by choosing which consumers to include in community experiments. Although they did not infer a consumption network, [53]’s microbial community assembly experiments found that the resulting community structure was highly variable at the species level, while highly predictable at the family level. Our theoretical results suggest a possible interpretation of these findings. When viewed at the species level, consumers do not differ enough to strongly affect the outcome of community assembly, suggesting that these species differences are below our analytical threshold and drift is the primary driver of abundance dynamics. On the other hand, consumer abundances converge to nearly deterministic outcomes at the family level, suggesting that between-family differences in resource profiles are well above our threshold.

## Supporting Information

**S1 Appendix**. Characterizing the spectrum of the deterministic model; Master equations for our stochastic model; Discussion of abundance distributions in the neutral and niche limits; Fitting outcomes using different cutoff values;

