## Supplementary material for "Emergent neutrality in consumer-resource dynamics": S1 Appendix

#### 1 Characterizing the Spectrum of the Deterministic Model

If a system of ordinary differential equations admits a fixed point, then the eigenvalues of its Jacobian evaluated at the fixed point tell us about the behavior of the system after a perturbation. The Jacobian in the system of differential equations in the main text evaluated at the abundances $r\vec{1}$ and $n\vec{1}$ is

$$L = \begin{bmatrix} -n[C\ \vec{1}]_{diag} & -rC \\ \epsilon n C^T & 0 \end{bmatrix} \tag{1}$$

where $\vec{1}$ denotes a vector of 1's and $[C\ \vec{1}]_{diag}$ denotes a diagonal matrix with entries given by the vector $C\ \vec{1}$. If the real parts of all the eigenvalues of the Jacobian $L$ are negative, then the system will return to equilibrium after a sufficiently small perturbation, and is therefore called locally stable. Previous work has shown that our model is inherently stable [1]. Here, we aim to analytically characterize the eigenvalue distribution of $L$ for both of our parametrizations of $C$, so we can estimate the rate at which the abundances return to equilibrium.

In Fig 1, we plot the spectrum of $L$ in the complex plane and see some characteristic features. There are two complex conjugate eigenvalues, one bulk of eigenvalues near zero and another bulk of eigenvalues centered at $-n\kappa$, where $\kappa$ is the average row sum of $C$. In the following sections, we derive approximations to the densities of these two real eigenvalue bulks, and predict the values of the complex conjugate eigenvalues.

In Fig 2, we plot the magnitude of the components of the eigenvectors corresponding to the two bulks of eigenvalues, and observe that in both the specialist and generalist cases, the eigenvalues near zero largely correspond to consumer directions, while those centered at $-n\kappa$ correspond to resource directions. After any kind of perturbation, we therefore expect resource abundances to quickly decay back to equilibrium, while consumer abundances will decay more slowly, because the eigenvalues which dictate their dynamics are small in magnitude. This time-scale separation recapitulates the classic expectation of fast resource dynamics relative to consumer dynamics, but doesn't alone tell us whether the system will be neutral-like, or not.

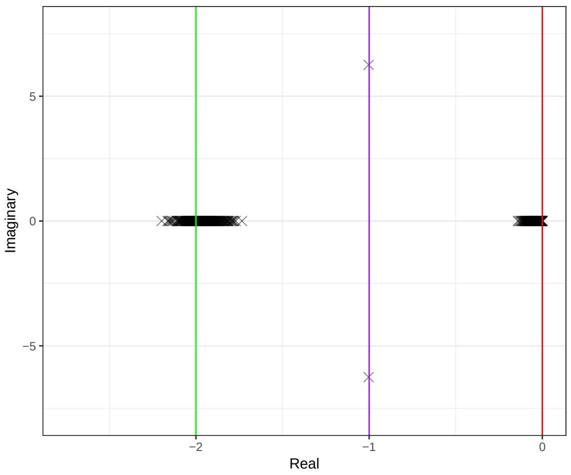


Figure 1: The spectrum of $L$ in the complex plane when $C$ is sampled from uniform distribution on $[0, 0.2]$, $K = S = 200$, $n = 1$, $r = 20$ and $\epsilon = 0.5$. The red line is $\Im(z) = 0$, the purple line is $\Im(z) = -\frac{1}{2}\kappa n$ and the green line is $\Im(z) = -\kappa n$.

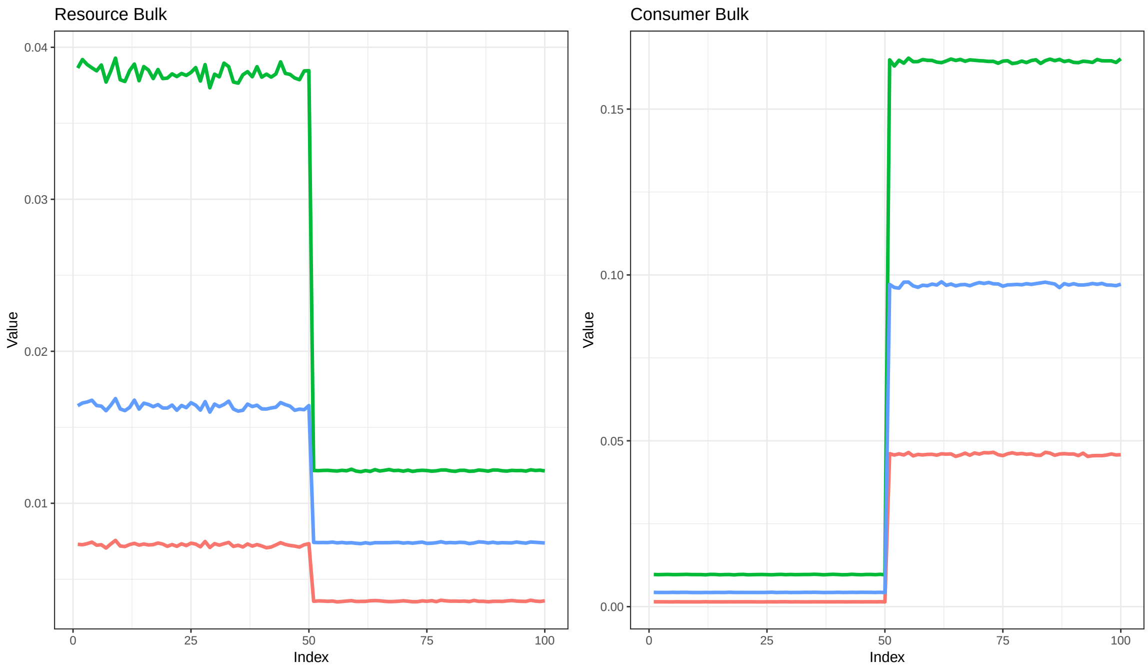


Figure 2: The first panel plots the first, second and third quartiles of the absolute values of the components of the eigenvectors associated to the eigenvalues centered at $-n\kappa$, while the second panel plots the same quantities for the eigenvectors associated to the eigenvalues near zero. The first quartile is in red, the median is in blue and the third quartile is in green. For these plots, the entries of the matrix $C$ are sampled from a uniform distribution on $[0.5, 1.5]$, $K = S = 50$, $s = r = 100$ and $\epsilon = 1$. The quartiles are computed using the absolute values of the components of the eigenvectors from $1000$ realizations of the matrix $C$.

#### 2 Master equation and abundance distributions in the neutral and niche limits

Let $Q_k(R_k|\vec{N},t)$ denote the probability that resource $k$ has abundance $R_k$ at time $t$ in our stochastic model. As mentioned, consumer dynamics are slow, so that $\vec{N}$ is approximately constant in time while resource dynamics play out. The master equation for $Q_k(R_k|\vec{N},t)$ is then

$$\begin{aligned}\frac{dQ_k(R_k|\vec{N},t)}{dt} &= \rho_k Q_k(R_k-1|\vec{N},t) + \left((R_k+1)\sum_{j=1}^{S} C_{kj}N_j\right) Q_k(R_k+1|\vec{N},t) \\ &\quad - \left(\rho_k + R_k\sum_{j=1}^{S} C_{kj}N_j\right) Q_k(R_k|\vec{N},t)\ .\end{aligned} \tag{2}$$

Because of our timescale separation, the sums in Eq 2 are constant, so the stationary distribution is simply a Poisson distribution with rate $\frac{\rho_k}{\sum_j C_{kj}N_j}$. In our simulations, we find that the distribution of a single resource throughout time, and also the distribution of resource abundances across the community, quickly converge to Poisson distributions, in agreement with Eq 2 (see Figs 3, 4).

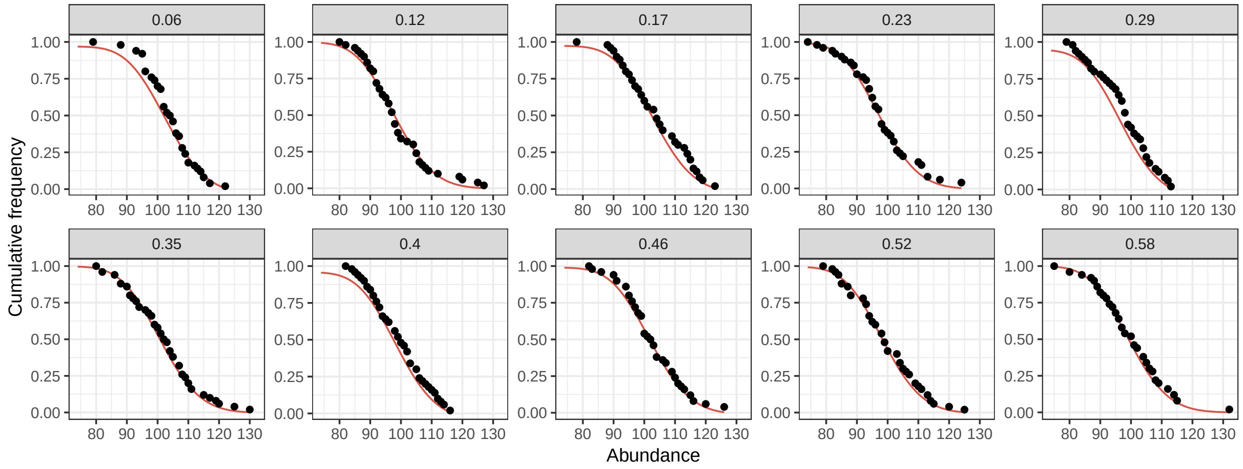


Figure 3: Abundance distribution of resources in the generalists scenario. Plots are faceted by the coefficient of variation of the consumption matrix. Red curves show the fitted Poisson distribution. The p-value of the Cramér-von Mises goodness-of-fit test is non-significant for all fits, reflecting successful Poisson fits to resource abundances across the board. Parameters: $n = 500$, $r = 100$, $S = K = 50$.

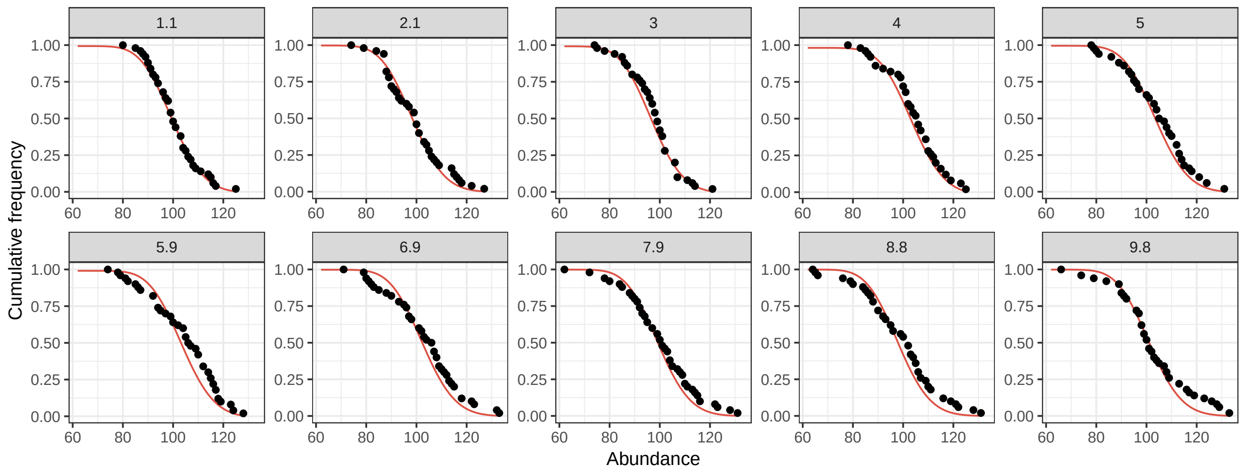


Figure 4: Abundance distribution of resources in the specialists scenario. Plots are faceted by the ratio of on-diagonal (preferred resource) to off-diagonal (all other resources) entries in the consumption matrix. Red curves show the fitted Poisson distribution. The p-value of the Cramér-von Mises goodness-of-fit test is non-significant for all fits, reflecting successful Poisson fits to resource abundances across the board. Parameters: $n = r = 100$, $S = K = 50$.

Next, we define $P_i(N_i|\vec{N}, t)$, the distribution of consumer abundance values $N_i$ over an ensemble of trajectories in our stochastic model. Since the resource abundances are tightly concentrated at their mean values, we assume that they are equal to their means, and we eliminate them from the master equation for consumer dynamics, leading to:

$$\begin{aligned}\frac{dP_i(N_i|\vec{N},t)}{dt} &= \epsilon(N_i-1)\left(\sum_{j=1}^{K}\frac{C^T_{ij}\rho_j}{C_{jj}(N_j-1)+\sum_{l\neq j}C_{jl}N_l}\right)P_i(N_i-1|\vec{N},t) \\ &\quad + \eta_i(N_i+1)P_i(N_i+1|\vec{N},t) - N_i\left(\eta_i+\epsilon\sum_{j=1}^{K}\frac{C^T_{ij}\rho_j}{\sum_l C_{jl}N_l}\right)P_i(N_i|\vec{N},t)\ .\end{aligned} \tag{3}$$

If the consumption coefficients are identical $C_{ij} = \mu$, then the sum $\sum_l C_{jl}N_l = \mu\sum_l N_l = \mu J$ is approximately proportional to the average of the stationary distribution for consumers, where $J$ is the total community size. If $J$ is exactly constant, then Eq 3 becomes the master equation for a neutral birth-death process, with a log-series stationary distribution. More concretely, if $C_{ij} = \mu$, then $\rho_i = \rho$ and $\eta_i = \eta$ for all $i$ and assuming detailed balance gives us the formula

$$P_i(N_i+1) = \frac{\epsilon}{\eta_i}\frac{N_i}{N_i+1}\sum_{j=1}^{K}\frac{C^T_{ij}\rho_j}{\sum_l C_{jl}N_l}P_i(N_i) = \frac{\epsilon S\rho}{J\eta}\frac{N_i}{N_i+1}P_i(N_i) \tag{4}$$

for all $i$. Recurring on Eq 4 gives a log-series stationary distribution with parameter $p = \frac{\epsilon S\rho}{J\eta}$ from the main text. So, when the consumption matrix has all identical coefficients, we expect a log-series stationary distribution as long as the fluctuations of community size $J$ are not prohibitively large at equilibrium. In simulations of the specialist scenario with $C_d/C_o$ varying between 1.1 and 18, the coefficient of variation of the total community size was low throughout our parameter values, never surpassing 0.03, and often as low as 0.01. Therefore, total abundance of consumers was largely invariant, justifying our assumption that $J$ is approximately constant. In this way, we see how neutral drift can plausibly emerge from a consumer-resource model when the differences in consumption preferences among consumers is small. As the coefficients $C_{ij}$ become more heterogeneous, consumer abundances are more strongly affected by specific combinations of consumers, and we expect drift to fail to predict the resulting abundance patterns, because it does not incorporate these relationships. For example, if consumers are completely specialized and $C$ is a diagonal matrix, then $\sum_l C_{jl}N_l = C_{jj}N_j$ is not a good estimate of the average value of the stationary distribution, since it is strongly affected by stochastic fluctuations in $N_i$ at equilibrium. In fact, the stationary distribution for Eq 3 is a Poisson distribution for each consumer in the limit consumers are pure specialists, reflecting the Poisson distribution of their specialized resources.

### 3 Simplifying the Jacobian

We make the approximation that the upper left hand block of our Jacobian is a diagonal matrix ($-n[C\ \vec{1}]_{diag} \approx -n\kappa I$). When $C$ is specialized, each row sum is precisely $(S-1)C_o + C_d$, so our approximation is exact. When $C$ is random, $\kappa = S\mu$ from the central limit theorem, but each row sum is not exactly constant. Let $\lambda$ be an eigenvalue of the Jacobian $L$. If $-(n\kappa+\lambda)I$ is invertible, then we can use block diagonal rules to find the eigenvalues of the Jacobian in Eq 1. On the other hand, if $\lambda = -n\kappa$, then $-(n\kappa+\lambda)I$ is not invertible and we want to find when $\lambda = -n\kappa$ is an eigenvalue of $L$. Let $\vec{v} \in \mathbb{C}^K$ and $\vec{w} \in \mathbb{C}^S$ be arbitrary vectors. We want to know if $L - \lambda I$ is singular, so suppose

$$\begin{bmatrix} 0 & -rC \\ \epsilon n C^T & -\lambda I \end{bmatrix} \begin{bmatrix} \vec{v} \\ \vec{w} \end{bmatrix} = \vec{0}\ . \tag{5}$$

Since $C$ is a $K \times S$ matrix and $K \geq S$, $C$ is injective in our parametrizations and hence Eq 5 implies $\vec{w} = \vec{0}$. ($C$ is injective if the equation $C\vec{w} = \vec{0}$ implies that $\vec{w} = \vec{0}$). However, $C^T$ is not injective when $K > S$, so $\lambda = -n\kappa$ is an eigenvalue of the Jacobian $L$ when $K > S$ with multiplicity $K - S$. Now that we understand the case when $\lambda = -n\kappa$, we may safely assume that $\lambda \neq -n\kappa$ and use block determinant rules to predict the remaining eigenvalues. We find that

$$\det[L - \lambda I] = \det[-\kappa n I - \lambda I] \det[\epsilon n r C^T((-\kappa n - \lambda)I)^{-1}C - \lambda I]\ . \tag{6}$$

If we compute the inverse and re-arrange some terms above, we get that the eigenvalues of $L$ satisfy

$$0 = \det[\epsilon n r C^T C - (-\kappa n - \lambda)\lambda I]\ . \tag{7}$$

Now, let $\omega$ be an eigenvalue of $C^T C$. Then,

$$\lambda = -\frac{1}{2}\left(n\kappa \pm \sqrt{(n\kappa)^2 - 4\epsilon n r \omega}\right) \tag{8}$$

are eigenvalues of $L$. So, to obtain the spectrum of $L$, we just need to determine spectrum of $C^T C$. Note that if $S > K$, then $C^T C$ has eigenvalues that are precisely zero and neutrality is assured.

### 4 Specialist Eigenvalues

In the specialist parametrization, the eigenvalues of $C^T C = C^2$ are simply the eigenvalues of $C$ squared. So, one eigenvalue is the square of the row sum $\kappa^2 = ((S-1)C_o + C_d)^2$ and the remaining eigenvalues are all equal to $(C_d - C_o)^2$. Therefore, we can use Eq 8 to predict the spectrum of $L$. The resource and consumer eigenvalues are given by

$$\lambda = -\frac{1}{2}\left(n\kappa \pm \sqrt{(n\kappa)^2 - 4\epsilon n r (C_d - C_o)^2}\right) \tag{9}$$

when the plus-minus is a plus and minus respectively. We use the formula for the consumer eigenvalues to derive the scaling relationship in the main text for the specialist parametrization. The outlying pair of eigenvalues is obtained by applying the transformation in Eq 8 to $\kappa^2$, and these eigenvalues will be complex when $4\epsilon r > n$.

For the generalist parametrization, the spectrum of $C^T C$ is more complex. In the following sections, we use random matrix theory to understand the spectrum of $L$ in the generalist case, and then calculate the eigenvalue relevant for predicting the drift threshold.

### 5 Generalist Outlying Eigenvalues

The Marchenko-Pastur law describes the spectrum of matrices whose entries are independent and identically distributed random variables with mean $0$ and variance $\sigma^2$. For our application to a consumer-resource model, we need to have $\mu > 0$ so that the entries of $C$ are rates of consumption. Sampling the entries of $C$ from a distribution with non-zero mean changes the spectrum of $C^T C$ by changing only one eigenvalue. Let's write $C = X + \mu 1_{K\times S}$, where $1_{K\times S}$ is a $K \times S$ matrix of $1$s and $X$ is a random matrix with mean $0$ and variance $\sigma^2$.

$$C^T C = (X^T + \mu 1_{S\times K})(X + \mu 1_{K\times S}) = X^T X + 2\mu X^T 1_{K\times S} + K\mu^2 1_{K\times S} \, . \tag{10}$$

We expect the row sums of this matrix to be about the same, and so one eigenvector of the matrix should be approximately $\vec{1}$ with an eigenvalue given by the nearly constant row sum. Therefore, the single outlying eigenvalue of a matrix following the Marchenko-Pastur law with non-zero mean should be approximated by $\omega_{\text{out}} \approx S(\sigma^2 + K\mu^2)$. Using the transformation in Eq 8, we find that the two outlying complex conjugate eigenvalues of the spectrum of $L$ are the image of the $\omega_{\text{out}}$. Fig 5 plots the observed and predicted imaginary parts of these outlying eigenvalues of $L$, while varying $\mu$ for different resource abundances $r$. We have chosen parameters for which $\omega_{\text{out}} = S(\sigma^2 + K\mu^2) > \frac{\kappa^2 n}{4\epsilon r}$ so that the real part of the outlying eigenvalues is simply $-\frac{1}{2}\kappa n$ and the imaginary part is given by the square root in Eq 8. Our predictions work well, and we now turn to the bulks of eigenvalues in the spectrum of $L$.

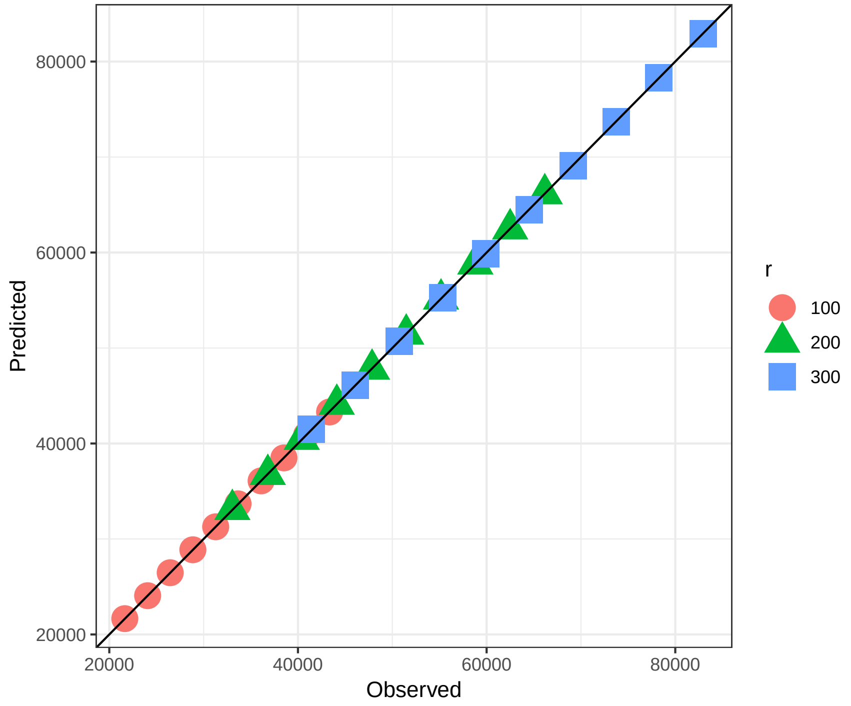


Figure 5: The plot shows the observed values of the imaginary part of the outlying eigenvalue of the Jacobian against our analytical prediction while we vary the average consumption strength $\mu$ between $0.5$ and $1$. The $C$ matrix is sampled from a uniform distribution on $[\mu - 0.5, \mu + 0.5]$, $K = S = 500$, $n = 100$, and $\epsilon = 1$. The color and shape of the points denote different $r$ values.

#### 6 Generalist Consumer Eigenvalues

Let $X$ be the matrix from the previous section. Let $\omega'_1 \leq \omega'_2 \leq ... \leq \omega'_K$ be the $K$ eigenvalues of $\frac{1}{K}X^TX$. Let's define the empirical spectral density of $\frac{1}{K}X^TX$ as

$$\nu_K(z) = \frac{1}{K}\sum_{i=1}^{K}\delta(\omega'_i - z) \; . \tag{11}$$

In 1967, Marchenko and Pastur proved that if $K, S \to \infty$ such that $S/K \to \gamma \in (0, \infty)$, then $\nu_K \to \nu$ where

$$d\nu(z) = \frac{1}{2\pi\sigma^2}\frac{\sqrt{(\gamma_+ - z)(z-\gamma_-)}}{\gamma z}\mathbf{1}_{[\gamma_-,\gamma_+]}dz \quad \text{with} \quad \gamma_\pm = \sigma^2(1\pm\sqrt{\gamma})^2 \tag{12}$$

when $\gamma \leq 1$ [2]. $\mathbf{1}_{[\gamma_-,\gamma_+]}$ is the indicator function on $[\gamma_-, \gamma_+]$. When $S > K$ so that $\gamma > 1$, $C^TC$ is not invertible, so there is at least one eigenvalue precisely equal to zero, and we recover the competitive exclusion principle – $S$ species require at least $S$ resources to stably coexist. The density in Eq 12 is called the Marchenko-Pastur law, and is controlled by $\sigma^2$, the variance in consumer preferences, and $\gamma$, the ratio between the number of consumers and resources. We are interested in the eigenvalues $\omega$ of $X^TX$ rather than $\frac{1}{K}X^TX$. Using Eq 12, we see that these eigenvalues are distributed in the interval $[K\gamma_-, K\gamma_+] = [K\sigma^2(1-\sqrt{\gamma})^2, K\sigma^2(1+\sqrt{\gamma})^2]$. So, we understand how the eigenvalues of $X^TX$ are distributed, and now we can use Eq 8 to predict the distribution of the consumer eigenvalues of the full Jacobian. The boundary of the support for the consumer eigenvalues is given by

$$\lambda_\pm = -\frac{\kappa n}{2}\left(1 - \sqrt{1 - \frac{4\epsilon r}{\kappa^2 n}K\sigma^2(1\pm\sqrt{\gamma})^2}\right) \approx -\epsilon r\frac{\sigma^2}{\gamma\mu}(1\pm\sqrt{\gamma})^2 \tag{13}$$

where we have expanded the square root to first order to obtain the approximation. $\lambda_+$ is the smallest (most negative) eigenvalue that governs local consumer dynamics, so it will correspond to the fastest relaxation time back to equilibrium, and hence it is the eigenvalue we use to derive our analytical prediction for when our stochastic consumer-resource system should appear neutral in the main text. In Fig 6, we plot the observed and predicted densities of the consumer eigenvalues of the Jacobian, as well as the predicted value of $\lambda_+$, for $\gamma = 1, \frac{1}{2}$. Our predictions work well in each case.

We should note here that we have really computed the eigenvalue distribution of the Jacobian where the upper left hand block is exactly $-n\kappa I$, which could be achieved exactly by, for example, imposing that every row sum of $C$ is the same constant. However, Fig 6 shows that the prediction for the eigenvalues of this simplified Jacobian still work well for the case of a non-constant upper left diagonal block.

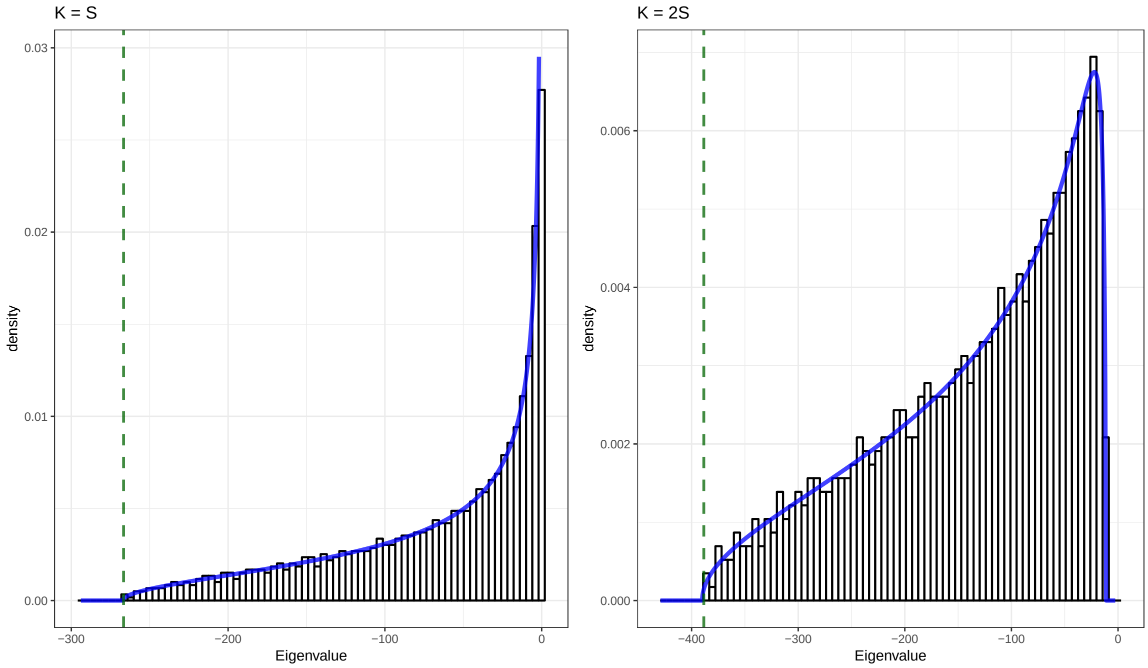


Figure 6: The two panels show histograms of the eigenvalues near zero in Fig 1 for different values of $K$ and $S$. For both panels, the entries of the $C$ matrix are sampled from a uniform distribution on $[0, 2]$, $s = 100$, $r = 200$ and $\epsilon = 1$. In the first panel, $K = S = 1500$, and in the second panel, $K = 2000$ and $S = 1000$. Our prediction for the density of eigenvalues from Eqs 8, 12 is the curve in blue. The dotted green line is our prediction for $\lambda_+$ – the smallest consumer eigenvalue and the eigenvalue we use to predict when the stochastic consumer-resource system should behave neutrally.

### 7 Generalist Resource Eigenvalues

In contrast to the eigenvalues near zero, the eigenvalues centered at $-n\kappa$ are not well described by the prediction assuming $-n[C\ \vec{1}]_{diag} \approx -n\kappa I$. Here, the heterogeneity in the values of the row sums becomes important for accurately predicting the distribution of resource eigenvalues. Although this is not mathematically justified, Eq 8 still gives us a reasonable approximation to the distribution of resource eigenvalues. We focus on Eq 8 when the $\pm$ is a plus:

$$\lambda = -\frac{1}{2}\left(n\kappa + n\kappa\sqrt{1 - \frac{4\epsilon r}{\kappa^2 n}K\omega'}\right) \approx -n\kappa + \frac{\epsilon r}{\kappa}K\omega'\ . \tag{14}$$

From the central limit theorem, we expect the row sums of $C$ to be normally distributed with mean $S\mu$ and variance $S\sigma^2$. If we take $\kappa$ to be a random variable with this distribution, we can use Eq 14 to find a new prediction for the resource eigenvalues. In fact, $n\kappa$ is much larger than the $\omega'$ term in Eq 14, so it is the distribution of $n\kappa$ that determines the predicted resource eigenvalues.

In Fig 7, we plot histograms of the observed resource eigenvalues with the prediction from Eq 14 without the $\omega$ term, and find that these predictions still work well. If we normalize the $C$ matrix so that all rows have the same sum and our approximation $-n[C\ \vec{1}]_{diag} \approx -n\kappa I$ is exact, then the resource eigenvalues are described by a Marchenko-Pastur law as predicted by Eq 14, with $K - S$ eigenvalues precisely equal to $-n\kappa$. So, the $\omega'$ term in Eq 14 can be important, but, for our parameter choices, the normally distributed row sums of $C$ seem to describe the resource eigenvalues reasonably well.

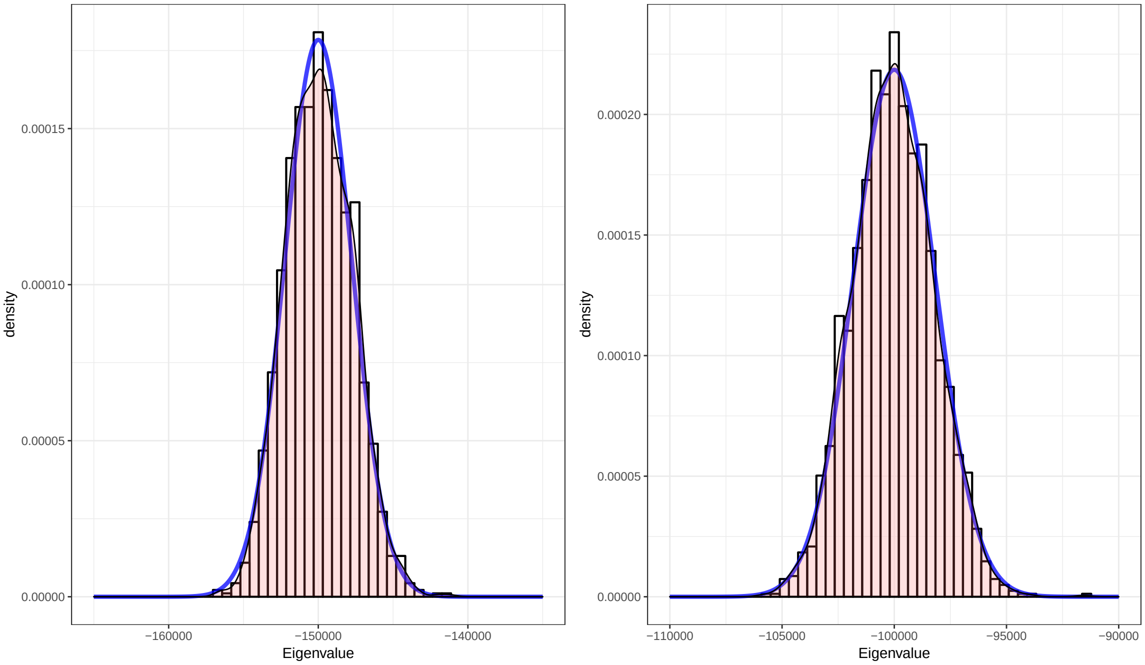


Figure 7: The two panels show histograms of the eigenvalues (with the corresponding densities in red) centered at $-n\kappa$ in Figure 1 for the same parameter values as in Figure 6. In the first panel, $K = S = 1500$ and, in the second panel, $K = 2000$ and $S = 1000$. We also plot in blue the prediction from Equation (14) while neglecting the corrections from the $\omega'$ term.

### 8 Different Cutoff Values

In the main text, we showed that as the consumption matrix departs from true neutrality, the log-series distribution becomes less likely to fit the species abundance distribution. We defined the point at which this probability falls below 50% as a threshold, which we compared with analytical predictions. While this 50% cutoff is arbitrary, our results are not sensitive to this choice. Specifically, our finding that $CV^{\text{threshold}}$ in the generalist scenario has a power law dependence on the species mean abundance with exponent $-0.5$ holds whether we define the cutoff at 50% or 5% or 90% (see Fig 8A, C). Similarly, our finding that the threshold ratio $C_d/C_o$ in the specialist scenario is linearly related to species richness holds across this wide range of choices for the cutoff (Fig 8B, D). This occurs because different cutoff choices lead to thresholds that differ only by a constant multiplicative factor, thus not affecting the functional dependence on $n$ and $S$.

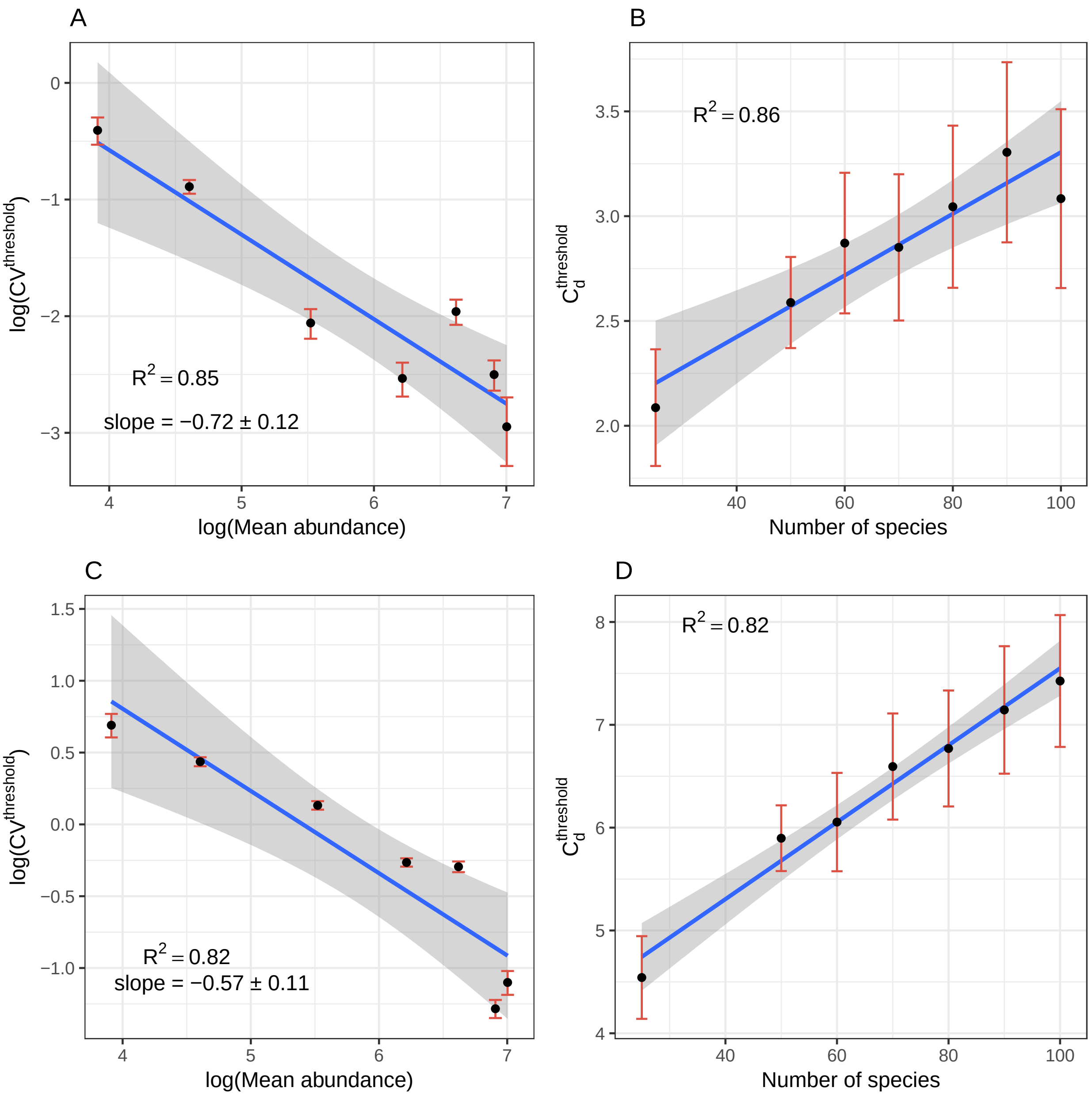


Figure 8: The functional dependence of the neutrality threshold on simulation parameters is robust to the choice of the probability cutoff. **A** and **B**: When we set the cutoff to 90% probability of a successful log-series fit, $CV^{\text{threshold}}$ in the generalist scenario (**A**) is still related to mean abundance via a power law with exponent close to $-0.5$. In the specialist scenario (**B**), the threshold ratio $C_d/C_o$ is still linearly related to the community richness. **C** and **D**: Similar results are obtained when setting the cutoff to 5% probability of successfully fitting the log-series. Compare with Fig. 2C, 2F in the main text. Notice how the slope in B (0.15$\pm$0.05) and D (0.03$\pm$) is different

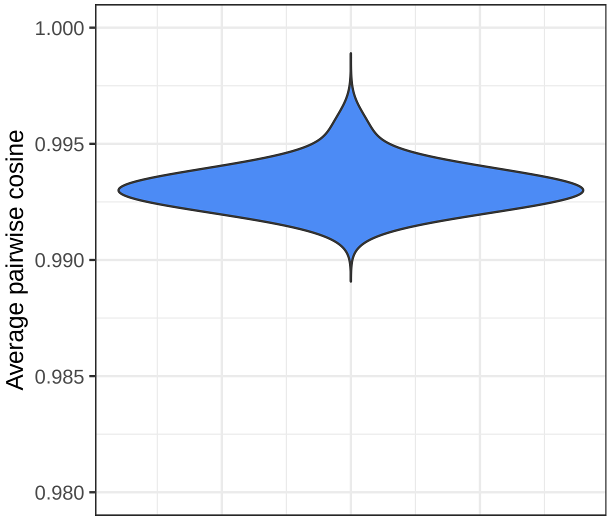


Figure 9: Distribution of mean pairwise cosines across an ensemble of neutral communities where every consumption event is recorded over an interval corresponding to the average species lifetime, and then used to estimate the consumption matrix. Namely, the estimated value for matrix entry is $\hat{C}_{ij} = \sum_k \frac{1}{R_{i,k}} \frac{1}{N_{j,k}}$, where $R_{i,k}$ and $N_{j,k}$ are the abundances of resource $i$ and species $j$ at the time of consumption event $k$, and the sum is over all consumption events involving this pair. From an ensemble of 878 such neutral communities with $S = K = 50$, $n = r = 100$, the 95% confidence interval for the mean pairwise cosines falls between 0.992 and 0.996. The average species lifetime given these parameters is 250 generations. If consumption events are observed in perpetuity, the estimated consumption matrix converges to the true (neutral) matrix, and the estimated mean pairwise cosine converges to 1.

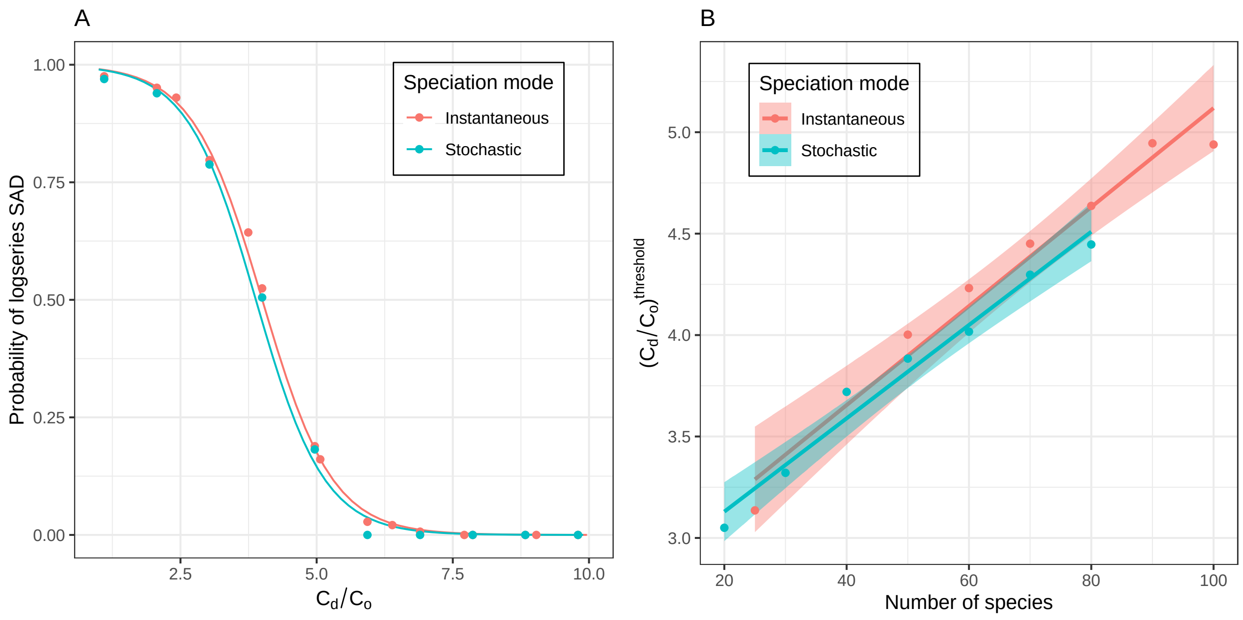


Figure 10: Comparison of RSA results between two speciation modes in the specialist scenario. **A**: Probability of a neutral fit plotted against the ratio $C_d/C_o$ of preferred to non-preferred resources, which quantifies the degree of niche differentiation between consumers (Number of species = 50). **B**: Linear behavior of the threshold against the number of species in the community. Results are statistically indistinguishable under either speciation mode (slopes: red: 0.02±0.002, blue: 0.02±0.002). Error bars are omitted for visual clarity.

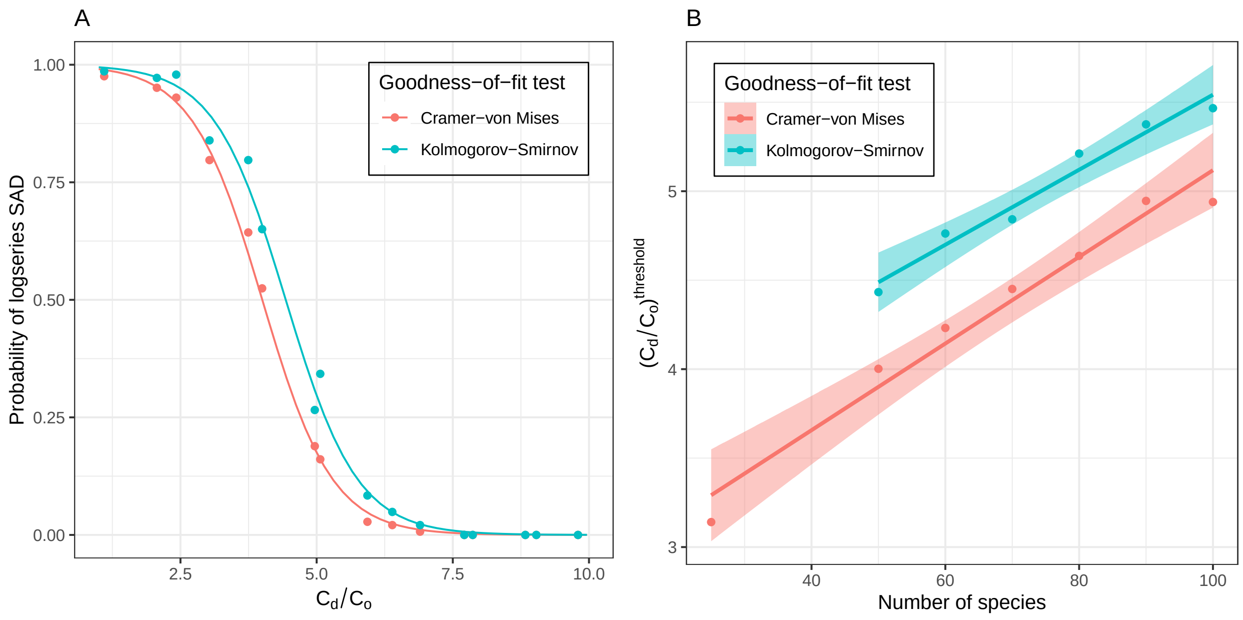


Figure 11: Comparison of RSA results between two different goodness-of-fit tests in the specialist scenario. **A**: Probability of a neutral fit plotted against the ratio $C_d/C_o$ of preferred to non-preferred resources, which quantifies the degree of niche differentiation between consumers. Both tests show a quick transition from high to low probability of a neutral RSA, with the threshold value of $C_d/C_o$, defined as the inflection point in the logistic regression, being slightly different between the two tests. (Number of species = 50). **B**: Both tests reveal linear behavior of the threshold against the number of species in the community. The threshold values are overall higher under the Kolmogorov-Smirnov test, indicating lower power to reject neutrality than the Cramér-von Mises test, however the linear scaling is identical, with indistinguishable slopes: C-vM: 0.02$\pm$0.002, K-S: 0.02$\pm$0.003. (Missing points for the K-S test are due to its being unable to reject neutrality in depauperate communities.)

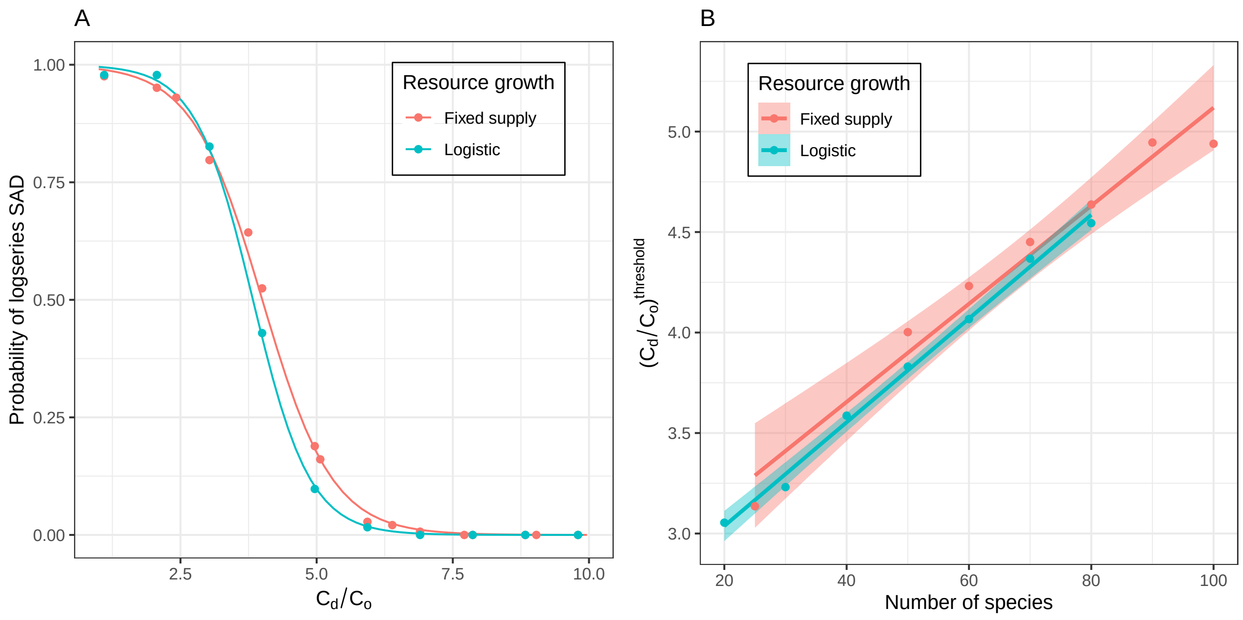


Figure 12: Comparison of consumer RSA results in the specialist scenario between two different systems of resource growth: fixed supply and logistic. Under the latter, the probability of a supply event to resource i is proportional to $r_0 R_i(1 - R_i/K)$ , where $r_0$ and $K$ are the intrinsic growth rate and carrying capacity of resources, respectively. **A**: Probability of a neutral fit plotted against the ratio $C_d/C_o$ of preferred to non-preferred resources, which quantifies the degree of niche differentiation between consumers. Both scenarios show a quick transition from high to low probability of a neutral RSA (Number of species = 50). **B**: The logistic scenario also shows linear behavior of the threshold against the number of species in the community ($R^2$ = 0.97). Slopes are similar between the two scenarios: fixed supply: 0.02±0.002, logistic supply: 0.03±0.0008.

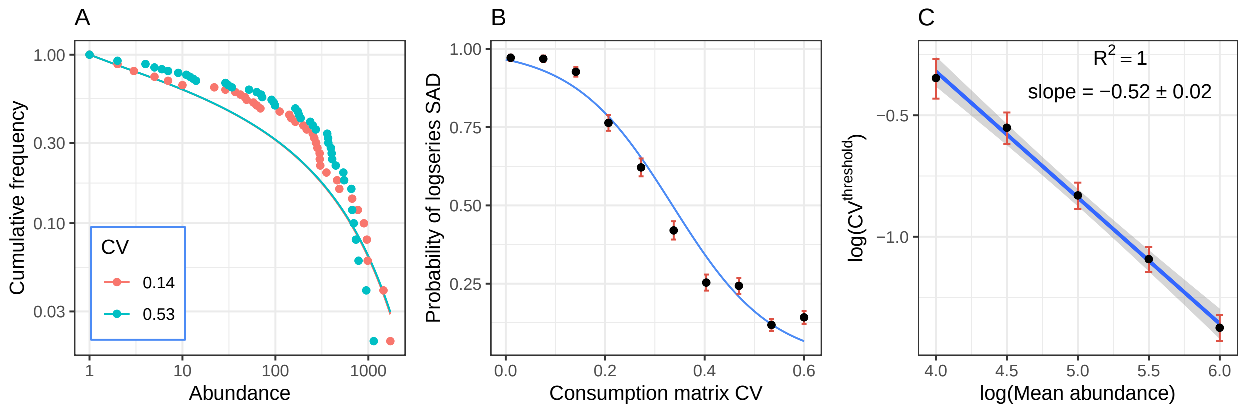


Figure 13: SAD results for the generalist scenario where the C matrix is drawn from a normal distribution. Results are analogous to those for C drawn from a uniform distribution presented in Figs 2A, B, C in the main text. **A**: The SAD fits a log-series (solid curve) when the coefficient of variation (CV) in the C matrix is sufficiently low (red points, Cramér-von Mises test p-value = 0.12), but not when the CV is sufficiently high (blue points, Cramér-von Mises test p-value = 0.005). **B**: The probability of a neutral fit is high for low coefficient of variation (CV) in the C matrix, and quickly decreases as the CV increases, with an inflection point at CV = 0.33. Points and error bars show the mean and standard error of the count of successful fits, out of an ensemble of 288 communities. Blue curve shows logistic regression. (Number of species = 50; Mean abundance = 245) **B**: The threshold CV, defined as the inflection point of the logistic regression, has a power-law dependence on the mean species abundance with fitted exponent -0.52$\pm$0.02, matching our prediction of -0.5.
